# Ultra-rare variants drive substantial *cis*-heritability of human gene expression

**DOI:** 10.1101/219238

**Authors:** Ryan D. Hernandez, Lawrence H. Uricchio, Kevin Hartman, Chun Ye, Andrew Dahl, Noah Zaitlen

## Abstract

The vast majority of human mutations have minor allele frequencies (MAF) under 1%, with the plurality observed only once (i.e., “singletons”). While Mendelian diseases are predominantly caused by rare alleles, their cumulative contribution to complex phenotypes remains largely unknown. We develop and rigorously validate an approach to jointly estimate the contribution of all alleles, including singletons, to phenotypic variation. We apply our approach to transcriptional regulation, an intermediate between genetic variation and complex disease. Using whole genome DNA and lymphoblastoid cell line RNA sequencing data from 360 European individuals, we conservatively estimate that singletons contribute ~25% of *cis*-heritability across genes (dwarfing the contributions of other frequencies). Strikingly, the majority (~76%) of singleton heritability derives from ultra-rare variants absent from thousands of additional samples. We develop a novel inference procedure to demonstrate that our results are consistent with rampant purifying selection shaping the regulatory architecture of most human genes.

## INTRODUCTION

The recent explosive growth of human populations has produced an abundance of genetic variants with minor allele frequencies (MAF) less than 1%^1^. While many rare variants underlying Mendelian diseases have been found^2^, their role in complex disease remains unknown^3–8^. Evolutionary models predict that the contribution of rare variants to complex disease is highly dependent on selection strength^9,10^, and that population growth can magnify their impact^10–12^. Recent methodological breakthroughs^13,14^ have enabled researchers to jointly estimate the independent contributions of low and high frequency alleles to complex traits, often demonstrating a large rare variant contribution likely driven by natural selection^5,15–18^. However, these studies excluded the rarest variants^15^ or included only well-imputed variants^5^ – this is a problematic limitation given that some plausible evolutionary models predict that the largest contributions to phenotypic variance could be from the rarest variants^9–11,19^. Directly querying the role of all variants with large-scale sequencing and sensitive statistical tests has the potential to reveal important sources of missing heritability, inform strategies to increase the success rate of association studies, and clarify how natural selection has shaped human phenotypes.

In this work, we develop, validate, and apply an approach for inferring the relative phenotypic contributions of all variants, from singletons to high frequency. We focus on the narrow-sense heritability (*h*^2^) of gene expression because a growing body of literature suggests that genetic variants primarily affect disease by modifying gene regulatory programs^20–23^, and recent examinations have identified significant rare variant effects on transcription^8^. To characterize the genetic architecture of gene expression, we analyze 360 unrelated individuals of European ancestry with paired whole genome DNA^24^ and RNA^25^ sequencing of lymphoblastoid cell lines (LCLs). We evaluate the robustness of our approach to genotyping errors, read mapping errors, population structure, rare variant stratification, and a wide range of possible genetic architectures.

## RESULTS

### Building and testing our model

Previous studies have sought to estimate the effect of rare alleles on trait variance, but have removed the rarest variants due to methodological challenges. Here, we developed a new method to solve this problem and validated our approach with an extensive set of simulations. Before analyzing real expression data, we performed a rigorous series of simulations to identify an approach for estimating heritability that is robust to possible confounding factors. In our simulations, we use real genotype data [all variants within 1 megabase (Mb) of the transcription start or end sites of genes] and generate gene expression phenotypes across individuals while varying the number of causal variants contributing to the phenotype (from 1 to 1,000), the distribution of effect sizes (including uniform, frequency-dependent, and an evolutionary based model), and the distribution of causal allele frequencies (ranging from predominantly rare to predominantly common; see Supplementary Table S1). In total, we simulated 440 different genotype-phenotype models that span the range of genetic architectures that likely underlie complex phenotypes such as gene expression, and analyzed each simulated dataset using multiple distinct methods. These include fitting a linear mixed model (LMM) via restricted maximum likelihood (REML^26,27^) and Haseman-Elston (H-E) regression, an alternative approach based on regressing phenotypic covariance on genotypic co-variance^26^ that is more robust in small samples (see Supplementary Figs S2, S3, and S17).

Similar to previous work^28^, we found that for many simulation settings, jointly analyzing all variants together can result in a substantial over- or underestimate of heritability (Fig. 1A, which shows results when true heritability is 0.2). One common solution is to partition sites by frequency^5,15,29^. We find that simply isolating rare (MAF<=1%) from common variants using two partitions and performing joint inference^15^ can improve the accuracy for most models. However, when there are many causal rare variants, the estimator remains upwardly biased. Partitioning alleles into five or more categories by MAF^5^ alleviates this problem. Remarkably, not only does the overall heritability bias decrease as the number of allele frequency categories increases, but Fig. 1B-F shows that the bias of the heritability for each MAF bin also decreases substantially across all models (see Supplementary Fig. S1). These simulations suggest that with our sample size, partitioning SNPs into 20 MAF bins results in the smallest bias in our estimate of total heritability (*h*^2^_total_) as well as the smallest bias for each bin across all simulated parameters (though see Supplemental Note 2.3 for further discussion of models that can induce bias). We note that further partitioning can improve results even further (e.g. Supplementary Fig. S6), but variance will likely increase unless prior knowledge about causal variation exists.

**Fig. 1.**
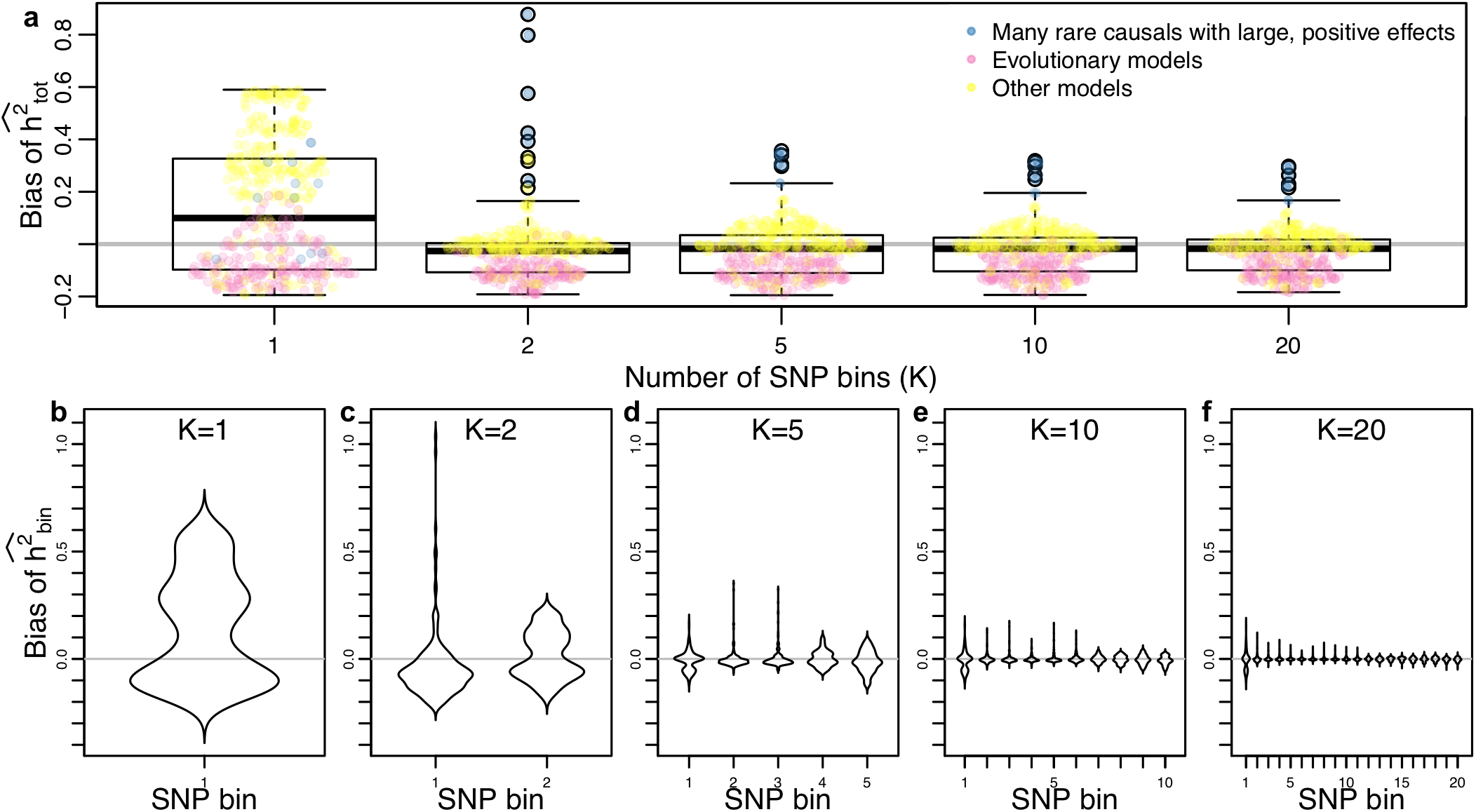
Simulation results showing that across a broad range of parameters, the accuracy of heritability inference improves as the number of SNP bins (partitioned by MAF) increases. (**A**) Mean bias of total heritability (inferred-true) for different numbers of SNP bins (**K**), where each point represents the mean of 500 simulations for different parameters, and a box plot summarizing the bias distribution across all parameters.(**B-F**) The distribution of average bias across simulated parameters for each SNP bin, showing that both mean and variance of the bias decrease as K increases.

When partitioning variants into multiple MAF bins, singletons are inevitably isolated into their own category. Intuitively, if some fraction of singletons are causal, then individuals with higher singleton load will be more likely to be phenotypic outliers (indeed, individuals with outlier expression patterns have been observed to have an enrichment of nearby rare variants^8^). It is therefore reasonable to ask what contribution singletons make to patterning phenotypic variation across a population. We investigated the theoretical properties of heritability estimation from singleton variants, and show analytically that when genotypic covariance is estimated using singletons alone, H-E regression is equivalent to regressing squared standardized phenotypes against singleton counts (see Supplementary Note 2.4).

A direct implication of our derivation is that H-E regression is unbiased unless singletons have large non-zero mean effect sizes (violating an explicit assumption of standard LMMs), which are the only simulation scenarios in Fig. 1A where heritability estimates remain upwardly biased (blue points). We develop an alternative approach that produces unbiased estimates of both heritability and mean effect size in all examined cases. Intuitively, the method (SingHer) conditions on total singleton count (per cis window) in order to (a) appropriately estimate total cis heritability and (b) partition singleton heritability into directional and random components (see Supplemental Note 2.4). However, because H-E regression is well understood and flexible, we recommend its use when mean effect sizes are near zero. For the data we analyze below, SingHer estimates that mean effect sizes are near zero, and we therefore proceed with H-E regression.

### Singletons drive the genetic architecture of human gene expression

In order to characterize the genetic architecture of human gene regulation, we partitioned the heritability of gene expression into 20 minor allele frequency (MAF) bins. We used *n*=360 unrelated individuals of European descent with both RNA sequencing data from GEUVADIS^25^ and whole genome sequencing data from 1000 Genomes Project (TGP)^24^. After extensive quality control to remove genes not expressed in LCLs, our data set includes 10,203 autosomal genes (see Supplementary Note 3.3). For each gene, we extracted all variants within 1Mb of the transcription start or end sites (corresponding to an average of 13,839 variants per gene, 35.2% are singletons); we do not consider *trans*-effects because of the small sample size (though we do analyze the effects of varying the window size in the Supplementary Fig. S12).

To control for possible non-normality, population structure, and batch effects, we quantile normalize expression values and include the first 10 principal components (PCs) from both the genetic and phenotypic data in all analyses, and present the average *h*^2^ estimate across genes in each MAF bin in Fig. 2A (blue curve). We find that *h*^2^is highest for the first MAF bin (singletons). However, using a novel trans-permutation procedure, we detected evidence for residual population stratification in low frequency (but not high frequency) SNPs that could not be accounted for using PCs (pink curve and see Supplementary Note 3.2, and note that differential population structure among common and rare variants is a documented, though understudied, phenomenon in human genetics^30^). We correct for this population stratification bias by subtracting the permutation-based estimate from the raw PC-corrected *h*^2^ estimate, shown in purple and henceforth indicated as *h*^2’^. We find that the plurality of *h*^2’^comes from singletons, but common variants also contribute a substantial amount towards *h*^2’^. Low and intermediate frequency SNPs make a minimal contribution to *h*^2’^. Note that this is a conservative correction because our *trans*-permutations capture both the effect of stratification and true *trans*-heritability.

**Figure 2.**
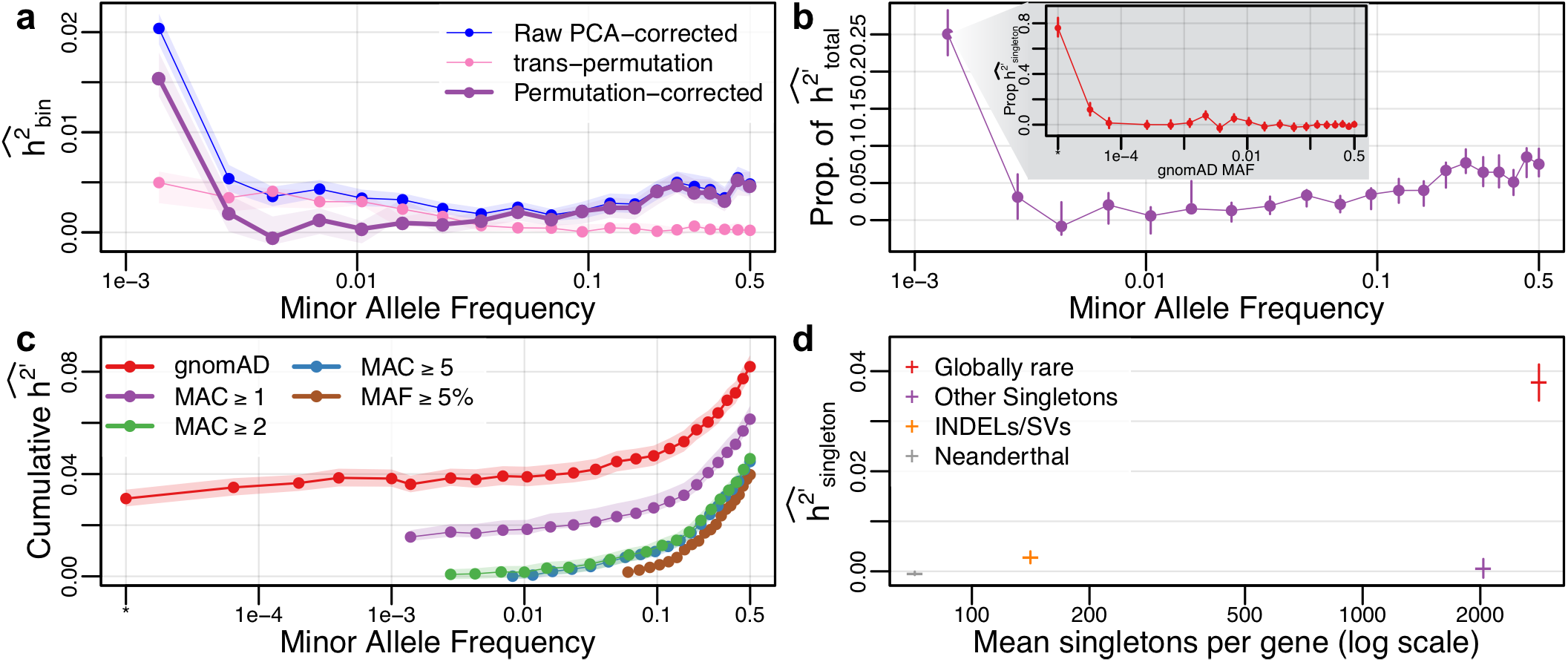
Rare variants (RVs) dominate the genetic architecture of human gene expression. (**A**) Average heritability estimates across genes, partitioned across minor allele frequency (MAF) bins (*h*^2’^, purple) after correcting for population structure using PCA (blue) and eliminating residual rare variant structure identified using a trans-permutation (pink). (**B**) The proportion of heritability attributed to each MAF bin. Singletons represent ~25% of the total inferred heritability, the vast majority of which is due to variants that are extremely rare in the population (inset, partitioning singletons in our data by the MAF observed in gnomAD, *n*>15k; singletons not reported in gnomAD are indicated by *). (**C**) Cumulative *h*^2’^ inferred as a function of MAF for different frequency filter thresholds (purple, green, blue, brown), and when singletons are partitioned by population MAF (based on gnomAD, red). Including all SNPs and partitioning singletons by population MAF (instead of observed MAF) results in a substantially increased level of *h*^2’^. (**D**) Globally rare singletons represent 56% of all singletons, but contribute 93% of *h*^2’^_singleton_. Rare INDELs and structural variants (SVs) also have enriched contributions to heritability (2.8% of singletons but 7.8% of *h*^2’^_singleton_). However, singletons inferred to derive from Neanderthal introgression or have gnomAD MAF≥10^−4^ make negligible contributions to *h*^2’^_singleton_. In all cases, confidence intervals/envelopes are based on the 95% quantile range of 1000 bootstrap simulations.

Fig. 2B shows the proportion of *h*^2’^ explained by each MAF bin, revealing that singletons represent ~25% of the total *h*^2’^, dominating the estimates from other MAF bins. Population genetic theory suggests that rare variants should only contribute a substantial fraction of *h*^2^ when causal variants are evolutionarily deleterious^9,10,12,31^. We therefore hypothesized that natural selection is the evolutionary force constraining the frequency of causal regulatory alleles. To test this hypothesis, we need to partition singletons according to the strength of selection acting upon them. However, since site-specific selection coefficients are generally not available for rare variants (except indirect measures such as functional prediction and evolutionary conservation, see Supplementary Fig. S20), we sorted our singletons by their population MAF, as inferred from a large, external database. We reasoned that some of the singletons in our dataset will be evolutionarily neutral with an intermediate population frequency, but the singletons that are most deleterious will almost always be constrained to low population frequency. We therefore partitioned singletons observed in our data by their MAF observed in the gnomAD dataset (representing high coverage whole genome sequencing on >15,000 individuals), and performed H-E inference of *h*^2’^ across 20 singleton bins based on their MAF observed in gnomAD. The inset in Fig. 2B shows that the vast majority (>90%) of singleton *h^2^*^’^ derives from variants that have gnomAD MAF<0.01%. This is strong evidence that natural selection constrains alleles with the largest effects on gene regulation to very low frequency. Note that we found that 31% of our singletons were not reported in gnomAD, but this subset of variants (indicated with MAF=“*” in Fig. 2B) nonetheless explains ~80% of *h*^2’^_singleton_. We confirm that the majority of this signal derives from true-positive singletons by analyzing a subset of 58 individuals with high coverage whole genome sequencing, and estimate that 88% of *h*^2’^_singleton_ derives from variants that validate (Supplementary Fig. S22). Previous work has shown that additionally partitioning common variants by LD resulted in minimal change after partitioning by MAF^5^.

Early studies of heritability filtered out SNPs with MAF<5% prior to their analysis^32^, and more recent studies only remove the rarest variants^5,15^. We show that the process of removing any SNPs based on MAF has a direct impact on the estimate of heritability. In Fig. 2C, we plot the cumulative *h*^2^ inferred as a function of MAF for different minor allele count (MAC) thresholds (averaged over all genes). We find that by adding progressively rarer variants to the analysis, there is a monotonic increase in the inferred heritability. Indeed, including all variants down to singletons (purple curve) increases inferred heritability by approximately 50% 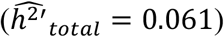 compared to the case when only common variants (MAF≥5%) are analyzed (brown curve, 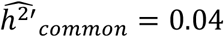), indicating that common variants are not able to tag heritability from lower frequency variants (see Supplementary Fig. S18 for additional analysis). However, not all singletons contribute equally to heritability, and pooling them together can deflate *h*^2’^ estimates (see Supplementary Fig. S6). Partitioning singletons into 6 bins based on their observed MAF in gnomAD (red curve) increases our 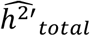 estimate to 0.082, and shows that nearly half of the total heritability (46.6%) is explained by the 27.6% of variants that are globally rare (with MAF_gnomAD_<0.1%). Note that *h*^2’^ represents the difference between raw *h*^2^ estimates and a conservative correction for population stratification, and hence these progressive increases in *h*^2^ cannot easily be explained by rare-variant stratification. We show analytically and with simulations that these results are consistent with a “singleton-LD” effect biasing estimates when rare variants are pooled together (see Supplementary Note 2.3.5), which previously has only been reported for common variants^5,28^.

To investigate the ability of rare variants to capture heritability of common variants (and vice-versa), we refit H-E regression removing MAF bins from rarest to most common (and vice-versa). We found that while rare variants could capture some of the heritability of more common variants, common variants could not capture the heritability derived from lower frequency variants (see Supplementary Fig. S18). This suggests that rare variants have not been indirectly captured in published heritability estimates through “synthetic association” tagging^33^.

Recent studies of gene expression variation in humans have suggested that one-quarter of Neanderthal-introgressed haplotypes have *cis*-regulatory effects^34^, and that expression outliers are enriched for having nearby rare structural variants (SVs) compared to non-outliers^8^. However, the overall contribution of these classes of variants to expression variation across genes remains unknown. We therefore performed H-E regression on four disjoint categories of singletons (Neanderthal-introgressed, indels/SVs, globally rare singletons, and other singletons), and found that globally rare singletons (i.e., singletons in our data that are also singletons across all 2,504 samples in 1000 Genomes Project^24^) contribute the vast majority (97%) of singleton heritability (Fig. 2D). Rare indels/SVs also have an enriched contribution to gene expression variation (representing 2.8% of singletons, but 6.8% of *h*^2^_singleton_), but Neanderthal-introgressed singletons and other singletons make a negligible contribution to *h*^2’^_singleton_.

### Genotype quality does not drive our inference of heritability

One possible confounding factor is the effect of genotyping error on heritability estimation^35^. If heritability is biased by genotyping error, and genotyping error also varies as a function of MAF, there could be differential bias across frequency bins when analyzing real data. We simulated a range of genotyping error models, and found that all investigated forms of genotyping error eroded efficiency of heritability estimation, but did not induce a detectable upward bias (Supplementary Fig. S5).

We also performed several analyses to examine possible confounding effects in these data (Supplementary Note 3.5). First, we ranked singletons by their reported genotype likelihood as reported for the individual carrying the singleton allele in TGP^24^, and partitioned them into four equal groups (quartiles). We then ran H-E regression with these four groups of singletons (along with 10 PCs). Strikingly, we find that only those singletons with high SNP quality contribute positively to our inference of heritability (see Supplementary Fig. S15). Second, since both the DNA and RNA sequencing are based on lymphoblastoid cell lines, it is conceivable that difficult-to-sequence regions of the genome could result in correlated errors that confound our inference. To test this, we restricted our analysis to regions of the genome passing the TGP Strict Mask^24^, and found that our inference of heritability was unchanged. We further ranked genes based on the number of exon bases passing the strict mask, and found no difference in the genetic architecture of genes having high versus low overlap with the Strict Mask (see Supplementary Fig. S16). Finally, a subset of n=58 samples were sequenced at high coverage by Complete Genomics Inc (CGI) as part of the 1000 Genomes Project^24^. We identified the singletons carried by these individuals, and partitioned them into four groups by cross classifying them as being present or absent in the CGI or gnomAD datasets. Running H-E regression on this subset of individuals shows that *h*^2’^_singleton_ is predominantly driven by singletons that replicate in the CGI data but are not reported gnomAD (consistent with Figure 2), and that singletons that are absent from CGI (and are therefore more likely to be false-positives) contribute negligibly to *h*^2’^_singleton_ (0.026 versus 0.005, respectively; Supplementary Fig. S22).

### Purifying selection drives the genetic architecture of gene expression

We found that rare variants are a major source of heritability of gene expression, which we hypothesized was due to purifying selection acting to constrain the frequencies of large-effect alleles. To test this hypothesis, we performed extensive simulations of human evolutionary history^36,37^, and developed a novel method to infer the parameters of an evolutionary model for complex traits (see Supplementary Note 4). Our three-parameter phenotype model extends a previously described model of the pleiotropy of causal variation^11^ (captured by *ρ*, where increasing values indicate higher correlations among expression effect sizes and the fitness effects acting on causal variants), and the scaling relationship between expression effect sizes and selection coefficients^9^ (*τ*, where increasing values indicate that distribution of effect sizes has a longer tail toward strong effects), to include the overall strength of selection (*ϕ*, a mixture parameter between strong and weak selection distributions, where *ϕ*=1 corresponds to strong selection). We inferred approximate posterior distributions for each of these parameters by rejection sampling^38^, which compares a set of informative summary statistics from genetic data simulated under a model of European demography^39^ and selection^40,41^ to the observed data (see Supplementary Note 4). Note that our inference procedure allows each parameter to vary across genes, but we only seek to infer the distribution of the average values of *ρ*, *τ*, and *ϕ* across genes because we do not have statistical power to infer *ρ* and *τ* for each gene. We rigorously evaluated the performance of this inference procedure with simulations, and found that we can infer *ρ* and *τ* with fairly high accuracy, but *ϕ* (while broadly unbiased) is less informative (Supplementary Fig. S23).

Applying this model to our data, we find that purifying selection has had a major impact on the genetic architecture of human gene expression, and that a range of previously explored evolutionary models can plausibly explain the empirical data. In Fig. 3A, we plot the posterior distributions of the mean values of *ϕ*, *ρ*, and *τ*, which suggest that on average: fitness effects acting on causal variants tend to follow the distribution inferred from conserved non-coding loci 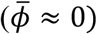, but selection is rampant in the sense that gene expression effect sizes are highly correlated with the fitness effects acting on causal variants. Fig. 3B shows that our data are consistent with a ridge of evolutionary scenarios that connect models in which causal alleles are highly modular (e.g. effect sizes are correlated with dampened fitness effects, as in the model of Eyre-Walker^9^, which assumes *ρ* = 1 with intermediate *τ*) and models with highly pleiotropic causal alleles and more extreme effect sizes (e.g., the Simons et al.^11^ model, which assumes *τ* = 1, but a more moderate *ρ*). This observation could only be identified using our integrated model, and suggests highly heterogeneous processes acting on individual genes. Interestingly, our parameter inference suggests that while mean *ρ*, *τ*, and *ϕ* can vary substantially among the best-fitting models, individual genes tend to have extreme values (i.e., either 0 or 1) for all three parameters (Fig. 3A). Fig. 3C shows the cumulative proportion of *h*^2^ as a function of MAF from 1,000 bootstrap draws from our posterior distribution, along with the cumulative proportion of *h*^2’^ inferred from our data. As compared to a neutral evolutionary model (pink), the posterior draws (grey, representing points along the ridge of evolutionary phenotype models show in Fig. 2B) are all highly concordant with our data.

**Figure 3.**
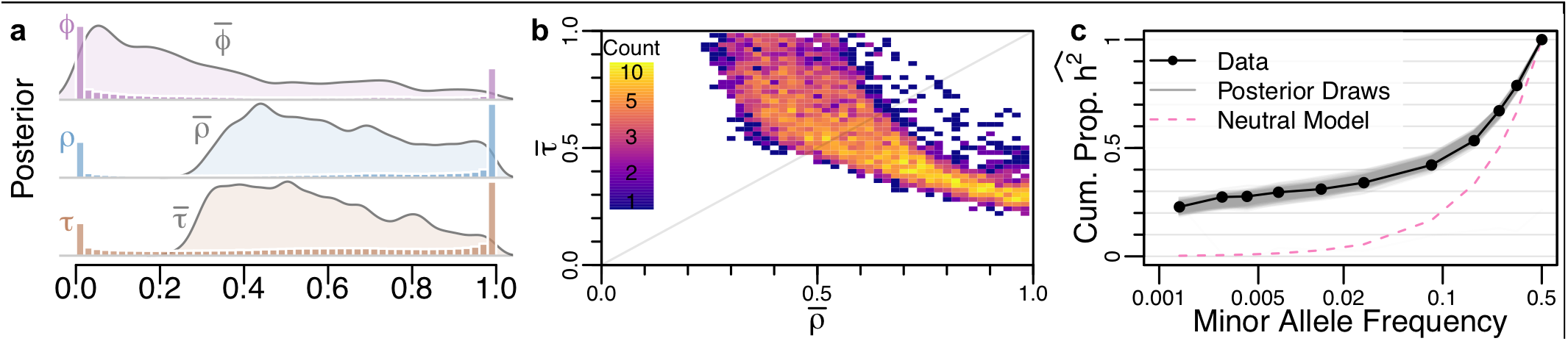
Rampant purifying selection drives the genetic architecture of gene expression. Our model infers the strength of purifying selection acting on causal variants (*ϕ*), the correlation between the fitness and the effect size of causal variants (*ρ*), and a scaling factor that transforms fitness into effect sizes (*τ*). (**A**) The posterior distribution of the mean of each parameter across genes (curves), as well as a histogram of the posterior parameter estimates for each gene. (**B**) The joint posterior distribution of the average r and t across genes shows an evolutionary tradeoff between the correlation and scaling of fitness and effect sizes. (**C**) The cumulative proportion of heritability inferred from the gene expression data (dots) compared to the expected patterns from 1000 draws from the posterior distribution (grey) and neutral expectation (pink).

## DISCUSSION

There is substantial interest in characterizing the genetic basis for complex traits to improve our understanding of human health and disease, and substantial resources are being spent to collect ever-larger cohorts to investigate the role of rare variants. Such studies will clarify what we have learned from our relatively small study of just 360 individuals. We developed, tested, and applied a novel technique for interrogating the role of rare variants in gene regulation using a relatively small cohort of *n*=360 individuals who had whole genome DNA and RNA sequencing performed on their derived lymphoblastoid cell lines. We estimate that the total narrow sense heritability of LCL gene expression is ~8.2%, and that the largest contributors to gene expression heritability are the rarest of variants in our data: singletons, where just one copy of the allele has been observed in our sample of 720 chromosomes (MAF=0.0014). Globally rare variants (MAF_gnomAD_ <0.01%) explain 90% of *h*^2^_singleton_, implying that many of these causal variants would remain singletons even if tens of thousands more samples were sequenced (and, concomitantly, many more singleton variants would emerge from the additional samples). This characteristic is best explained by rampant purifying selection, where most *cis*-acting regulatory variants are deleterious. We developed a rejection-sampling algorithm to infer parameters of an evolutionary model that are consistent with these data, and found that for ~2/3 of genes, effect sizes of *cis*-regulatory variants are highly correlated with how deleterious the fitness effects are on causal variants. Further, for the majority of genes, the fitness effects are more consistent with broadly defined conserved non-coding regions of the genome^40^ than the strongly selected nonsynonymous^41^ or ultra-conserved regions of the genome^42^. However, while these parameters allow us to generate simulated data consistent with our observations, they remain *post hoc* parametric models that do not necessarily represent a generative model of how the genetic architecture of *cis*-regulatory variation evolved, and do not incorporate potentially important contributions from other modes of natural selection (such as positive selection or balancing selection, which may be rare but can have substantial impact on gene expression when they act^43^).

Our estimate of total *cis*-heritability is slightly larger than the previous estimates of *h*_*cis*_^2^=0.057 and *h*_*cis*_^2^=0.055 in blood and adipose respectively^44^, but lower than recent twin-based estimates of overall narrow-sense heritability *h*^2^=0.26, 0.21, and 0.16 in adipose, LCLs, and skin respectively^45^ as well overall broad-sense heritability *H*^2^=0.38 and 0.32 for LCLs and whole blood^46^. It is therefore plausible that rare variants account for some “missing heritability” in human gene expression, but differences in population, tissue, and/or technology could also explain some of these patterns, and there could also be differences between the genetic architecture of *cis*-regulation and *trans*-regulation.

A concurrent examination of rare variant heritability via an allele specific expression approach^47^ reports a lower, but still substantial, contribution to heritability from rare variation. However, there are fundamental differences between our analyses that likely contribute to the difference in estimates. First, their work examines a much narrower window around genes. This will lead to differences if selection has acted differently in promoters compared to more distal regulatory regions^48^ (cf. Supplementary Fig S12). Second, their work uses a smaller sample size and so their definition of rare is less stringent than ours. Finally, they do not reclassify rare variants according to external reference panels, which greatly increased our estimates of rare variant heritability.

While it might at first seem logical to genotype some (or all) of these singletons in a larger panel of individuals to statistically identify the causal ones, our analysis uncovered a major challenge with this approach: our results can only be explained if the causal alleles driving heritability are evolutionarily deleterious, with effect sizes often scaling with the strength of selection acting on them. This means that the alleles that have the greatest impact on gene expression are likely to be extremely rare in the broader population, and are unlikely to exist in more than a few unrelated individuals in any given population. This is consistent with a recent finding that a large fraction of individuals with outlier expression for a gene also tend to have a globally rare variant in the vicinity^8^. We push this result further to quantify the overall impact that rare variants have on gene expression across a population. Indeed, we find that globally rare variants are the predominant source of heritability for gene expression. Our analysis shows that 90% of the singleton heritability derives from alleles that are either not reported or have MAF<0.01% in the *n*>15,000 samples in gnomAD. We therefore conclude that identifying causal alleles for transcriptional variation will likely require the incorporation of new biological information, possibly including large-scale experimental testing of singleton variants to improve functional predictions.

Our results suggest that one cannot capture the heritability of rare or low frequency alleles by analyzing additional common alleles. This implies that “synthetic associations”^33,49^ are uncommon for gene expression data. A broader consequence is that, when rare variants matter, approaches that rely on genotyping large samples followed by imputing missing genotypes from reference populations may not successfully reconstruct the true impact of rare variants (especially when the reference panel is smaller than the test sample). This is because both genotyping and imputation require the variant to be present at a reasonable frequency in the reference population, which is highly unlikely for strongly deleterious alleles (indeed, we found that ~80% of our singleton heritability was attributable to variants not reported in the >15,000 samples in gnomAD). Instead, WGS of large cohorts may be necessary (though the actual sample size required will depend on several factors that have not yet been elucidated).

As the number of samples with detailed phenotype data and WGS data increases, it will be possible to apply the approach we have developed here to characterize the genetic architecture of additional complex traits. Indeed, in a recent WGS study of height and BMI, we found that rare variants compose essentially the entirety of “missing heritability” for these traits^50^. By integrating such methods with functional genomic data, we may also learn more about the biology of causal variants, which could enable improved identification of clinically actionable variants in some cases. However, it is not clear that risk prediction from genomic data for most diseases will be feasible for otherwise healthy individuals with limited family history information. Population genetic theory tells us that rare variants will only be a significant source of heritability when causal alleles are evolutionarily deleterious. But the biology of human health and disease is complex. While not all human diseases will themselves impart a strong fitness effect, extensive pleiotropy resulting from tightly interconnected networks of interacting proteins experiencing cell-specific regulatory mechanisms could. Indeed, under the omnigenic model of disease, variants that affect any one of these components could contribute to an individual’s risk for any disease involving any down-stream pathway^23^.

We developed an approach to examine the heritability of singleton variants, and the results have important implications for future genetic studies. We rigorously evaluated the performance of our inference procedure using extensive simulations and multiple types of permutations (see Supplementary Fig. S21). While we employed several approaches to test for the presence of confounders from population structure, genotyping/mapping error, and cell line artifacts, there may be other unknown confounders that have biased the results of this study. We conservatively used quantile normalization on the expression phenotypes to enforce normality, and this often reduces the overall heritability estimates (see Supplementary Figs. S7 and S11) by diminishing the impact of outliers^8,51^. There are several other contributors to broad sense heritability that we have not attempted to model and may also account for some of the heritability estimated in family-based studies, such as gene-gene interactions, gene-environment interactions, and other non-additive components.

## SUPPLEMENTARY NOTE

Supplementary Note includes Supplemental Methods and Procedures, 27 figures, and three tables.

## AUTHOR CONTRIBUTIONS

R.D.H and N.Z. conceived of and designed the study. L.H.U. and A.D. developed methods. R.D.H., L.H.U., K.H., C.Y., A.D., N.Z. contributed to data analysis or simulations. R.D.H. and N.Z. wrote the manuscript. All authors read and approved the manuscript.

## ACKNOWLEDGMENTS

We thank Hyun Min Kang, Adam Auton, Sasha Gusev, and two anonymous reviewers for discussions about possible confounders that improved our analysis; members of the Pritchard lab for comments on rejection sampling; Jeffrey Barret and Konrad Karczewski for peer review of our preprint; Raul Torres for assistance with data analysis; Jeffrey Wall for assistance with Neanderthal-introgressed alleles; and Aria Hernandez for discussions on figure colors. Research reported in this publication was supported by National Human Genome Research Institute of the National Institutes of Health under award number R01HG007644 to RDH and K25HL121295. LHU was supported by IRACDA NIGMS grant K12GM088033, KH was supported by a Gilliam Fellowship, and AD was supported by U01HG009080 and R01HG006399.

