## Supplemental Information for "Ultra-rare variants drive substantial *cis*-heritability of human gene expression"

### 1 Table of Contents


**Ultra-rare variants drive substantial cis-heritability of human gene expression** **1**

**2 Partitioning heritability by minor allele frequency** **3**
*2.1 Overview of model and method* *3*
*2.2 Haseman-Elston (H-E) regression* *3*
*2.3 Evaluating the model with lots of simulations* *4*
2.3.1 Which simulated models cause bias? 5
2.3.2 Comparing H-E and REML. 5
2.3.3 H-E is robust to the fraction of variants that are causal 6
2.3.4 Simulating genotype error 6
2.3.5 The "Singleton-LD" effect – Simulations with globally rare causal variants vs random singletons. 7
*2.4 Further investigation of singleton heritability with theory.* *9*
2.4.1 H-E is the same as regressing singleton count on squared phenotype. 10
2.4.2 Proving H-E is the same as regressing singleton count on squared phenotype. 11
2.4.3 H-E for Singleton LMMs is broadly unbiased 12
2.4.4 H-E is biased when the mean singleton effect is nonzero 13
2.4.5 Singleton LMMs can identify mean effect sizes 14
*2.5 Software availability.* *15*

**3 GEUVADIS data set and QC** **15**
*3.1 RNA-seq processing.* *15*
*3.2 PCA and Permutations: controlling for population structure and batch effects.* *16*
*3.3 Identifying high quality genes.* *16*
*3.4 Robustness to model parameters.* *18*
3.4.1 Number of SNP bins and Quantile Normalization. 18
3.4.2 Window Size. 19
3.4.3 Number of PCs. 20
*3.5 Genotype quality and genome mapability.* *20*
*3.6 Comparing H-E and REML* *21*
*3.7 Common variants tag a negligible amount of $h^2$ from rare variants* *21*
*3.8 Heritability as a function of evolutionary conservation.* *22*
*3.9 Population structure makes a negligible contribution to singleton heritability.* *22*
*3.10 Analyzing a subset of individuals with high coverage whole genome sequencing* *22*
*3.11 Heritability enrichment across MAF.* *23*
*3.12 Exploring population covariates as an alternative to PCA.* *23*
*3.13 Constructing bootstrap confidence intervals.* *23*

**4 Inferring posterior distributions of evolutionary parameters** **24**
*4.1 Overview of rejection sampling pipeline* *24*
*4.2 Evolutionary model of complex phenotypes* *25*
*4.3 Forward in time-simulations* *25*
*4.4 Simulated phenotypes* *26*
*4.5 H-E on simulated expression values and genetic data* *26*
*4.6 Parameter inference* *26*
*4.7 Validation* *27*
*4.8 Sufficiency of summary statistics for parameter estimation* *27*
*4.9 Predicting the population-wide frequency of causal singletons.* *28*

**5** **30**

### 2 Partitioning heritability by minor allele frequency

#### 2.1 Overview of model and method

Given genotypes at $M$ SNPs over $N$ individuals we consider additive phenotypic models such that the phenotype of individual $i$ is: $y_i = \sum_{j=1}^{M} g_{ij}\beta_j + \epsilon_i$; $\epsilon_i \sim N(0, \sigma_e^2)$, where $g_{ij}$ is the genotype of individual $i$ at SNP $j$, $\beta_j$ is the effect size of SNP $j$, and $\epsilon_i$ is the residual, i.i.d. normally distributed noise of individual $i$. We partition the SNPs into $K$ disjoint sets determined by the minor allele frequency (MAF) of each SNP and wish to estimate the contribution of SNPs in the $k^{\text{th}}$ set to the heritability of $y$: $h_k^2 = \sigma_k^2 / \sigma_y^2$; where $\sigma_k^2$ is the genetic variance contributed by all of the SNPs in the $k^{th}$ partition, $\sigma_g^2 = \sum_{k=1}^{K} \sigma_k^2$ is the total genetic variance, and $\sigma_y^2 = \sigma_g^2 + \sigma_e^2$ is the total phenotypic variance, assumed equal to 1 going forward.

Recent work from several groups has examined this problem in the context of whole-genome sequencing, exome-sequencing, and imputation of large population cohorts[1–3]. Two of these studies partitioned SNPs into two or five MAF bins and jointly estimated the heritability in each bin using a linear mixed model fit by GREML. We similarly consider partitioning SNPs into bins by MAF, but investigate a larger range of bins from $K$=1 to 20. For computational and statistical reasons, we estimated heritability using Haseman-Elston (H-E) regression, which we describe below.

#### 2.2 Haseman-Elston (H-E) regression

To infer the heritability of gene expression levels across individuals, we primarily rely on H-E regression[4]. The premise of H-E regression is that heritability can be estimated by the correlation between the phenotypic covariance across individuals and the genotypic covariance across individuals. In practice, for a single gene, we estimate the phenotypic covariance ($P$) as the upper triangle of the outer product of quantile-normalized $\log_2$(FPKM) across our sample. For each of the K partitions, we estimate genotypic covariance with the upper triangle of a kinship matrix generated from all SNPs in the partition. Given a standardized genotype matrix of SNPs in the $k^{\text{th}}$ partition ($G_k$, with $N$ rows and $M_k$ columns, where each column has mean 0 and unit variance), the $k^{\text{th}}$ kinship matrix is $R_k = G_k G_k{}' / M_k$. H-E regression is then performed using the lm() function in R:

$$P \sim R_1 + \cdots + R_K$$

Specifically, the regression is ordinary least squares applied to the (vectorized) strict upper triangles of these matrices (which, for $N$ individuals has $\binom{N}{2}$ elements). In H-E regression, the effect size for the $k^{\text{th}}$ term represents the genetic variance explained by the $k^{\text{th}}$ SNP partition ($\beta_k = \sigma_k^2$), with the total genetic variance explained by all SNPs given by $\sigma_g^2 = \sum_{k=1}^{K} \sigma_k^2$. In the absence of other genetic contributions to phenotypic variation, heritability is equal to the total additive genetic variance explained by SNPs, $h^2 = \sigma_g^2$. Therefore, in most instances in this paper we simply refer to the genetic variance explained as heritability.

Since our H-E analysis is just regression, it is straightforward to add covariates to control for additional sources of variation as we describe here. We typically include the first 10 principal components (PCs) generated from our genome-wide genotype matrix as well as the first 10 PCs generated from our transcriptome-wide expression matrix (though we show below that the number of PCs included does not qualitatively impact our results). Formally, we include the $j^{\text{th}}$ PC (or an arbitrary numerical covariate) by adding the upper triangle of the PC's (or covariate's) outer product with itself to our symbolic regression equation above. Note that we were unable to obtain unbiased results by regressing out PCs or covariates from the phenotype vector and genotype matrix, but further research would be necessary to understand this behavior.

#### 2.3 Evaluating the model with lots of simulations

To evaluate the properties of our estimation method, we first examined several genetic architectures for a hypothetical phenotype. In most of our simulations, we use real genotype data by randomly sampling genes from the genome and extracting all genetic variants within 1Mb of their transcription start and end sites (though see below for effects of changing the window size). Our simulation parameters for the genetic architecture are described in **Table S1**. A phenotype is simulated by first choosing $h^2$ and $r$, the number of causal variants. We then chose one of three ways to determine the causal variants: 1) randomly sample $r$ variants near the gene; 2) choose $r_{rare}$ proportion of variants with MAF $\leq f$ and the remaining $r\times(1-r_{rare})$ with MAF $> f$; or 3) choose $r$ variants from the Uricchio et al. evolutionary phenotype model[5]. Under the Uricchio et al model, phenotype effect sizes are determined by ρ and τ. For the other two methods of choosing causal variants, we implement three different models for simulating effect sizes: 1) constant effect sizes; 2) i.i.d. N(0,1); and 3) the frequency dependent model of Wu et al.[6]: $0.4\times\log_{10}(\text{MAF})$. Finally, effect sizes are scaled to give the chosen heritability, $h^2$.

Varying all of the parameters in **Table S1** along with our methods for simulating effect sizes results in 2,200 different parameter combinations that were evaluated in our study (each with 500 independently simulated datasets). Each set of simulated phenotypes was analyzed multiple ways. In particular, we varied $K$, the number of SNP bins, and minMAC, the minimum allele count for a SNP to be included in the analysis. **Figure 1** (main text) shows the particular case when $h^2$=0.2, minMAC=1, and K is varied. **Figure S1** shows all values of $h^2$ and minMAC.

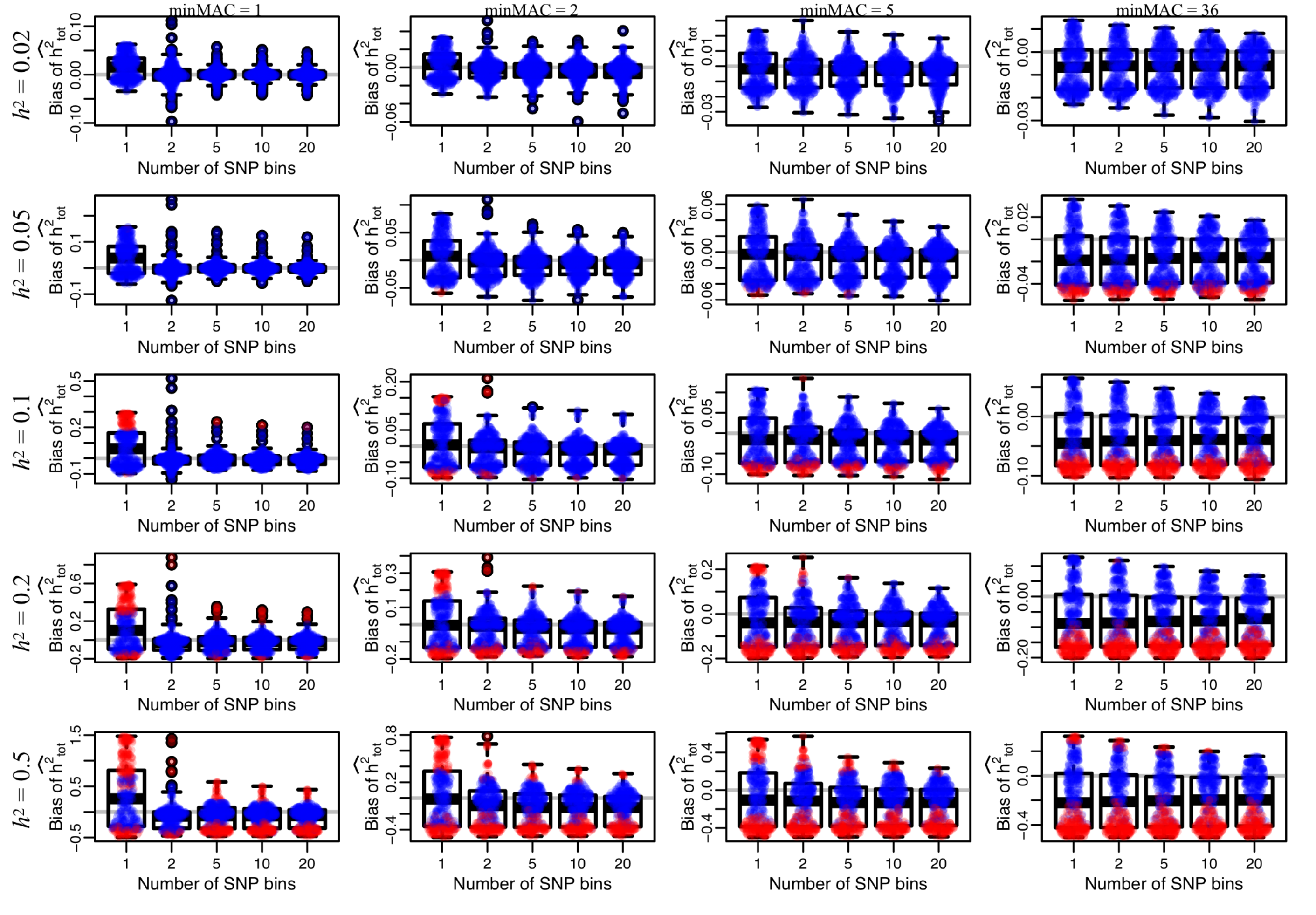


**Figure S1.** Simulation results showing bias of our H-E estimate of total $h^2$ across all parameters, analogous to Fig 1A (main text), varying the level of $h^2$ (rows) and the minimum minor allele count (minMAC) threshold for inclusion (columns; including all variants down to singletons on the left, and excluding all variants with MAF<5% on the right). Points are red if the central 90% empirical interquantile range across 500 simulations did not encompass the true $h^2$ and blue otherwise. For many simulated models, choosing K=1 or excluding low frequency variants leads to substantial bias.

**Table S1**. Parameters for simulating genetic architecture.

| Parameter | Description | Simulated values tested |
|---|---|---|
| $h^2$ | Total heritability | 0.02, 0.05, 0.1, 0.2, 0.5 |
| $\boldsymbol{r}$ | Number of causal variants | 1, 10, 100, 1000 |
| $\boldsymbol{r_{rare}}$ | Fraction of causal variants that are "rare" | 0.01, 0.05, 0.1, 0.5, 1.0 |
| $\boldsymbol{f}$ | Frequency threshold for rare variants | 0.01, 0.05, 0.1 |
| ρ | Effect size-fitness effect correlation | 0, 0.5, 0.8, 0.9, 0.95, 1.0 |
| τ | Effect size-fitness effect scaling factor | 0.5, 0.8, 1.0, 1.5 |

##### 2.3.1 Which simulated models cause bias?

We find that, in general, analyzing all SNPs in a single bin or excluding rare or low frequency variants from the analysis leads to substantial bias across many simulated models. These analytical approaches are therefore not robust to variation in the underlying genetic architecture (which for a real trait we may know little about). Jointly analyzing all SNPs partitioned into 20 disjoint bins by MAF results in the most consistent estimate of $h^2$ across the bevy of simulated genetic architectures that we have analyzed.

However, when $h^2$ is large (e.g. $h^2$=0.5), we find that even with K=20 and minMAC=1, several models exhibit downward bias (underestimating $h^2$), and a few that exhibit upward bias (overestimating $h^2$). The models that overestimate $h^2$ all follow particular pattern: $r$=1000, $r_{rare}$=1, $f$=0.01 or 0.05, and all effect sizes are in the same direction (either fixed or frequency dependent). In words, upward bias occurs when there are many causal variants (all of which are rare) with a substantially biased direction of effect. Such models seem unlikely to predominate complex traits in humans.

Models that result in underestimates of $h^2$ even when K=20 and minMAC=1 also share consistent patterns. A few models that result in underestimates have very few causal variants ($r$=1 or 10) that are all chosen to be rare ($r_{rare}$ = 1.0 and $f$=0.01 or 0.05). The vast majority of models resulting in underestimates stem from the Uricchio et al evolutionary model, primarily when $N_c$ is small (either 1 or 10) or when both ρ and τ are near 1. **Figure 1A** (main text) shows points colored by category of simulation, and demonstrates that there are very few counter examples to these broad characteristics.

Note that all simulations in **Figure S1** include quantile normalization of the simulated phenotype (just as we do with the real expression data, to be conservative). Below we investigate the role of quantile normalization and show that it can lead to substantial underestimates of $h^2$.

##### 2.3.2 Comparing H-E and REML.

The common approach for estimating $h^2$ from unrelated individuals in the literature is to use REstricted Maximum Likelihood (REML) to fit a linear mixed model (LMM), often as implemented in GCTA[7]. We implemented the Average-Information (AI) algorithm to fit LMMs in R to facilitate direct comparisons with our simulations[8]. **Figure S2** shows the mean relative bias [(inferred – true)/true] across all simulations where REML converged using typical criteria. When K=1, almost all simulations converge (**Figure S3**), and we see REML slightly out-performs H-E. Unfortunately, AI does not converge for many simulated models, particularly with large $K$, and as the number of simulations that converge with the AI algorithm decreases, the mean estimate of $h^2$ becomes un-stable. In **Figure S3**, we show the number of simulations for which we obtained an estimate for $h^2$ from H-E regression (black) versus REML (blue). We find that when K=10 or 20, AI rarely converges for any simulated model, while H-E has an analytic solution that can always be calculated. We did not experiment with adding covariates, but our real data analysis below suggests that AI would fail to converge with even smaller $K$ if they were included.

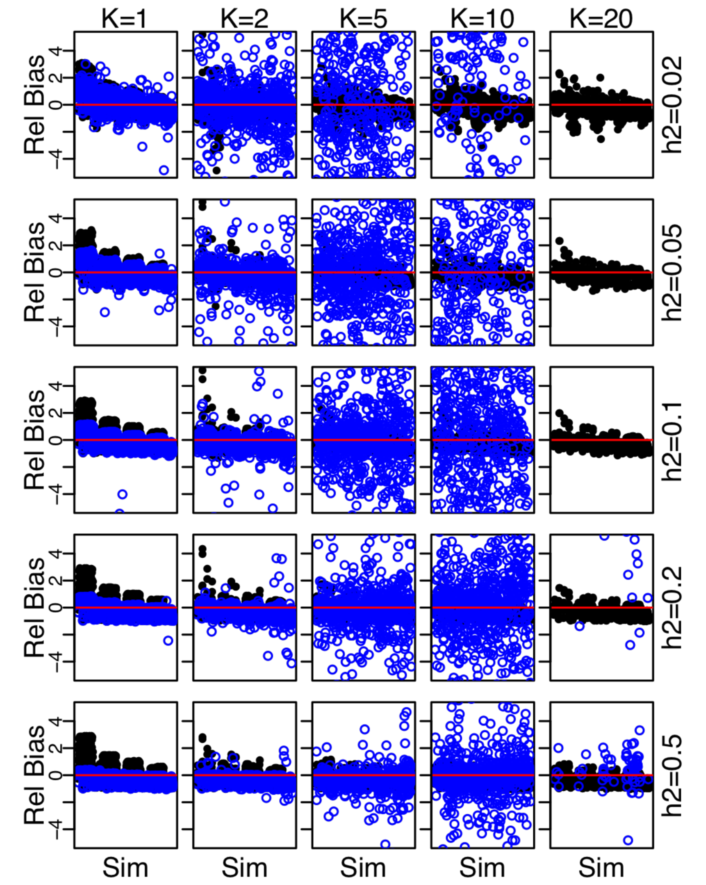


**Figure S2.** Mean relative bias [(inferred-true)/true] for each set of simulated parameters shown in Fig S4 for H-E (black) and LMM (blue).

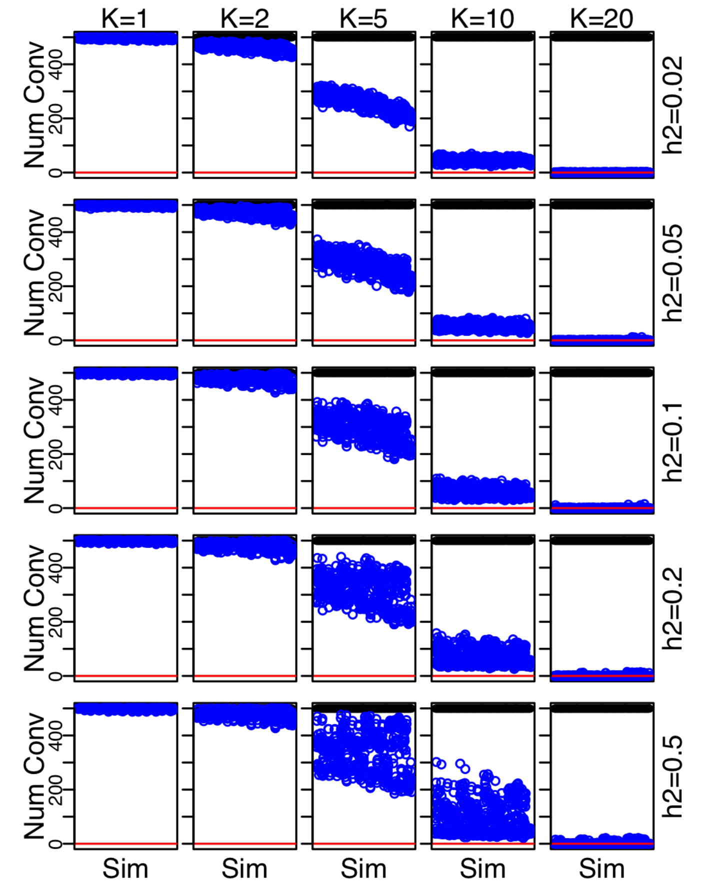


**Figure S3.** The number of simulations that converge for H-E (black) and LMM (blue), where each point is a set of simulated parameters. As the number of SNP bins increases, LMM struggles to converge but H-E regression does not.

##### 2.3.3 H-E is robust to the fraction of variants that are causal

When analyzing data, one does not necessarily know whether a trait is highly polygenic or not. We therefore varied the fraction of SNPs in a window that are causal from very small (0.01%) to very high (50%). We varied the true level of $h^2$, and ran simulations to test whether our inference of $h^2$ depends on the fraction of variants that are causal. We find that our inference procedure is robust to the fraction of variants that are causal (though some underestimate of $h^2$ occurs when a very large fraction of variants are causal and true heritability is very large), but the standard error is larger when only a small fraction of variants are causal (**Figure S4**).

##### 2.3.4 Simulating genotype error

To understand the impact of genotyping error on our inference about $h^2$, we performed the following simulations. Suppose there are separate error rates for false-positives and false-negatives. There is substantial evidence showing that these two rates should not be the same in short read data from the TGP[9–11]. In our simulations, only alternate alleles can be causal, so any reference allele has no effect on phenotype. A false-negative arises when an individual carries one causal allele at a site, but their genotype is called as homozygous reference; or alternatively the individual carries two causal alleles but are called as either heterozygous or homozygous reference. Such scenarios can arise particularly frequently with low coverage data and conservative genotype callers. It is also possible for an individual to truly be homozygous reference (i.e. carry no causal alleles at a site), but genotyped incorrectly to carry one or two causal allele(s). False-positives are thought to be less common than false-negatives genome-wide in the TGP data, but we do not know where causal variants lie and therefore simulate across a much

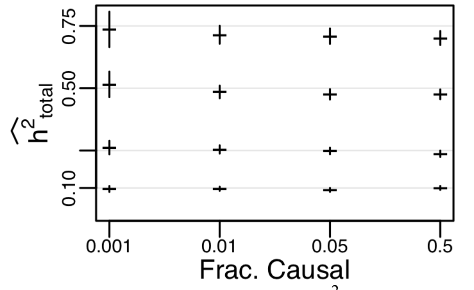


**Figure S4.** We varied the true $h^2$ (grey lines) and the fraction of variants that are causal. We find that H-E is largely robust (mean±2se).

larger range than expected in the data. We varied both the false-negative ("missing") rate and false-positive ("errAdd") rates, and performed 5000 simulations for each parameter combination. We simulate data under a model similar to the one we infer, where ρ=0.9, τ=0.5, ϕ=1, and the true $h^2$=0.2 with 100 causal variants. We randomly sample 5000 genes and use all the variants within 1Mb of the TSS or TES. For each simulation, we estimate $h^2$ using H-E regression with K=20 bins. In **Figure S5**, we show the mean ± 2 s.e. for $h^2_{total}$ across all 5000 simulations. Adding errors as either false-positives or false-negatives at causal loci can result in a slight underestimate of $h^2_{total}$, but in these cases all confidence intervals around the mean overlap the true value.

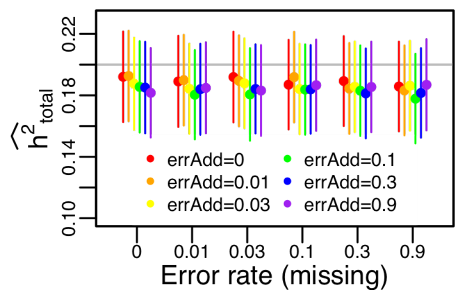


**Figure S5. Simulating genotyping error does not induce bias.** We simulate separate error rates for missing causal genotypes (e.g. low coverage induced false-negative; x-axis) and for adding false-positive genotypes at causal loci (colors). We generated 5,000 simulations for each parameter combination, and plot the mean±2se.

##### 2.3.5 The "Singleton-LD" effect – Simulations with globally rare causal variants vs random singletons.

One defining feature of our analysis is the investigation into the contribution of singletons to heritability. To demonstrate that our inference is capable of robustly inferring heritability due to singletons, we performed an additional set of simulations. In contrast to the simulations above, which were performed to guide the analytical approach for our real data, the simulations we discuss here were performed after observing the pattern in **Figure 2B** (main text). We constructed a simulation to test whether the inferred increase in heritability from globally rare alleles (relative to all singletons) is plausible. For the 10k genes included in our analysis, we extracted all singletons and noted their global minor allele frequency (gMAF) in the TGP-wide dataset (*n*=2505, which allows us to avoid missing data issues with the gnomAD data). We then specified a fraction of all singleton variants to be causal (fracCausal=0.001, 0.01, 0.05, 0.5) and sampled two sets of causal variants. In one set, fracCausal singletons were chosen from those variants that were singletons in the TGP-wide dataset (i.e., globally rare), and in the other set fracCausal singletons were chosen randomly from all singletons in our sample of n=360 individuals. Using these two sets of causal variants, we generated two phenotypes vectors for each gene (one based on the globally rare singletons, and one based on the randomly chosen singletons), where causal variants have i.i.d. Gaussian effect sizes, and we varied $h^2$ among {0.1, 0.25, 0.5, 0.75}. We then inferred $h^2$ for both types of phenotypes from all SNPs (rare and common, even though only singletons are causal) in two ways. First, we partitioned all SNPs into K=20 bins based on the observed frequency in our *n*=360 samples (as with data analysis shown in green in **Figure 2**, main text). Second, we partitioned SNPs into K=20 bins based on their frequency across all samples in TGP (*n*=2505; analogous to the red curve in **Figure 2C**, main text).

In **Figure S6**, we show the results of these four analyses. We find that when causal variants are chosen to be globally rare, and we partition all SNPs by global allele frequency, our inference is unbiased (red points). However, pooling all singletons together can result in a substantial underestimate of $h^2$ (blue points). This recapitulates our observation in **Figure 2C** (main text), which shows that partitioning by gnomAD MAF (red curve) results in a higher estimate of $h^2$ compared to partitioning variants by the MAF observed in our *n*=360 dataset (purple curve). We refer to this effect as "Singleton-LD". To our knowledge, this is the first time $h^2$ has been partitioned by allele frequency in an external dataset, and is therefore the first time this effect has been documented. Our finding strongly suggests that this approach should be considered in future studies.

One challenge that arises is the use of quantile normalization. In **Figure S7** we show that under many circumstances, performing quantile normalization results in an underestimate of $h^2$, but the pattern remains the same: when globally rare variants are causal but all singletons are analyzed in a single group, our estimate of $h^2$ is even more biased downward. To be conservative, we apply quantile normalization in the observed data anyway, since there is potential concern for non-normality in the observed gene expression patterns (though we show below there are no qualitative effects of quantile normalization).

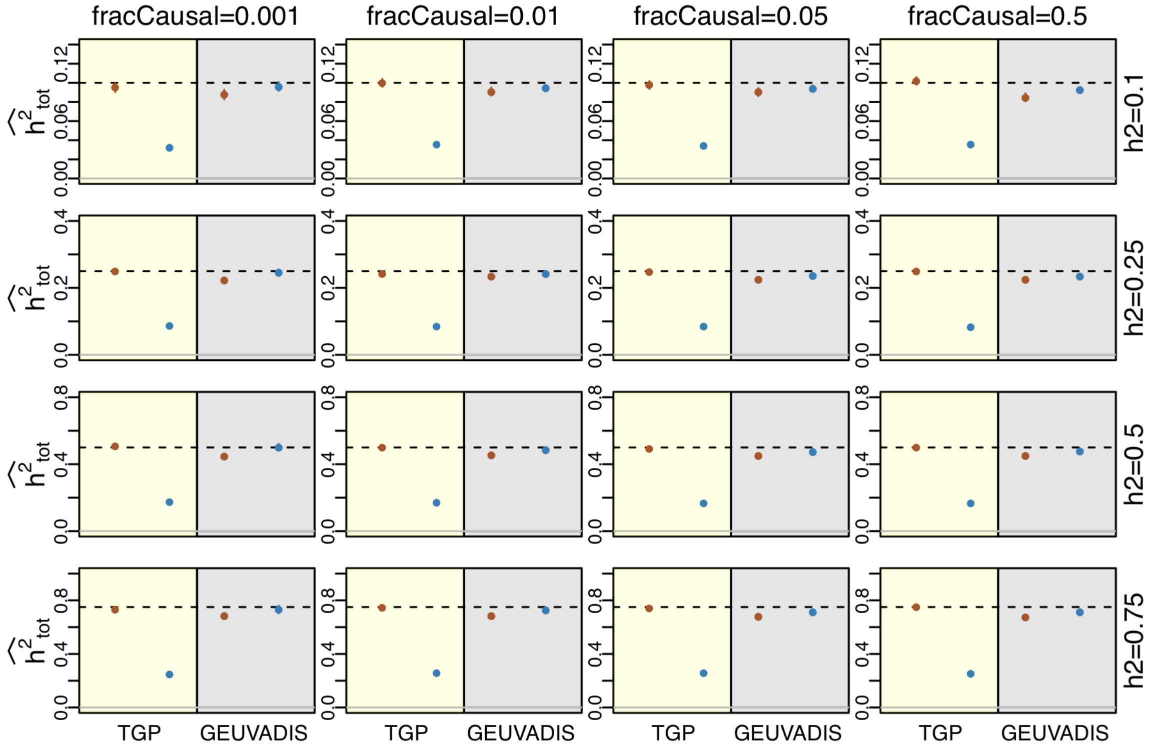


**Figure S6.** Singleton simulations without quantile normalization is unbiased. Inference of total $h^2$ when analyzing all SNPs but all causal variants are singletons for different levels of $h^2$ (rows) and different fraction of singletons that are causal (fracCausal; columns). Points in yellow shading indicate that causal singletons are chosen to be globally rare, while grey shading indicates causal singletons are chosen randomly from all singletons. Red points indicate partitioning all alleles by TGP-wide allele frequencies while blue points indicated partitioning by GEUVADIS frequency. When causal variants are globally rare, pooling all singletons together results in a substantial under estimate of $h^2$, but when causal singleton variants are randomly chosen, partitioning by TGP frequency does not increase bias very much. Points are means across all 10k genes and whiskers (often too small to see) represent 2SE.

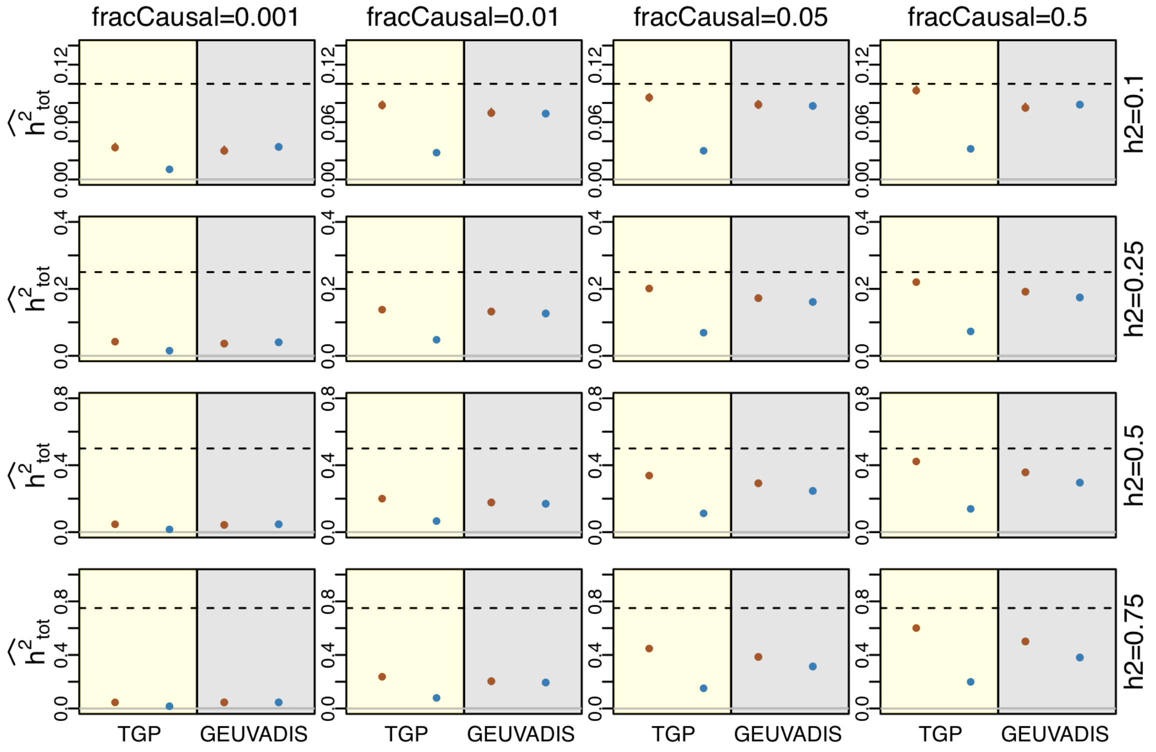


**Figure S7.** Simulated data are the same as Fig S2, but phenotypes are quantile normalized (as with the real data). Inference of $h^2$ is substantially biased downward. However, quantile normalization can avoid possible problems when observed data are not normally distributed.

#### 2.4 Further investigation of singleton heritability with theory.

In the case of $N$ individuals and $M$ SNPs, we assume the standard linear mixed model (LMM) for an $N$×1 phenotype vector $y \in \mathrm{R}^{\mathrm{N}\times 1}$ and an $N$×$M$ SNP genotype matrix $G \in \{0, 1, 2\}^{N\times M}$:

$$y = G\beta + \varepsilon$$

$\beta_j \sim N\left(0, \frac{1}{M}\sigma_g^2\right)$ independent for all SNPs $j$

$\varepsilon_i \sim N(0,\ \sigma_e^2)$ independent for all individuals $i$ (and independent of $\beta$)

If we define $u := G\beta$ to be the genetic contribution to the trait, then heritability is given by

$$h^2 := \frac{Var(u)}{Var(y)}$$

We additionally assume that $G$ consists of only singletons. This means that each column $j$ of $G$ will have $N$-1 "zeros" and 1 "one", and so has mean and variance 1/$N$. And $u$ simplifies:

$$u_i := (G\ \beta)_i = \sum_{j=1}^{M} G_{ij}\beta_j \triangleq \sum_{j:G_{ij}=1} N\left(0, \frac{1}{M}\sigma_g^2\right) \sim N(0,\ x_i\ \sigma_g^2)$$

where $\triangleq$ informally means equal in distribution and $x_i$ is defined as the fraction of singletons carried by individual $i$:

$$x_i := \frac{\#\text{ singletons for person } i}{\#\text{ singletons total}} = \frac{\sum_j G_{ij}}{M}$$

Although LMMs generally model between-sample phenotypic and genetic similarity, there is zero genetic similarity across singletons, so the $u_i$ are independent. Because the $\varepsilon_i$ are also independent, the joint model on the phenotype vector $y$ simplifies to independent marginal models on each observation:

$y_i \sim N(0,\ x_i\sigma_g^2 + \sigma_e^2)$ independently for all individuals $i$ (1)

Similarly, the heritability is simple to evaluate:

$$h^2 = \frac{E\big(Var(u|x)\big) + Var\big(E(u|x)\big)}{E\big(Var(y|x)\big) + Var\big(E(y|x)\big)} = \frac{E\left(x\ \sigma_g^2\right)}{E\left(x\ \sigma_g^2 + \sigma_e^2\right)} = \frac{\frac{1}{N}\sigma_g^2}{\frac{1}{N}\sigma_g^2 + \square_e^2}$$

(The inside expectation/variance operators, conditional on $x$, are over the random variables $\beta$ and $\varepsilon$, while the outside operators are over the empirical distribution of $x$.)

The factor $1/N$ accounts for the fact we defined $\beta$ on the scale of the genotypes $G$ rather than on the scale of the standardized genotypes as is customary. The normalization involves dividing the columns of $G$ by their standard deviation—for singletons, this is $1/\sqrt{N}$ for every column, and so the $\beta$ on the conventional scale have variance $\frac{1}{NM}\sigma_g^2$.

##### 2.4.1 H-E is the same as regressing singleton count on squared phenotype.

The Haseman-Elston (H-E) heritability estimate is constructed from $K \in \mathrm{R}^{\mathrm{N}\times\mathrm{N}}$, an inter-sample genetic similarity matrix, and the phenotypes $y \in \mathrm{R}^{\mathrm{N}\times 1}$. We take the standard approach and define K as the sample covariance of the column-demeaned genotype matrix:

$$K = \frac{N}{M} P_1^{\perp} G G^T P_1^{\perp}$$

This uses the demeaning projection $P_1^{\perp} := I_N - \frac{1}{N} 1_N 1_N^T$, where $I_N$ is the identity matrix and $1_N$ is a vector of 1s; this matrix is useful because for any $z$, $z' := P_1^{\perp} z$ is the demeaned version of $z$, and is central to the behavior of H-E with singletons. The $N$ factor is equivalent to standardizing columns of $G$ to variance 1 (for general, non-singleton $G$, the columns will have different variances and the ordinary column-wise scaling cannot be replaced by a single scalar multiplication, as we have done here with $N$).

Again, because $G$ contains singletons, $G_{i,}G_{j,}^T$ is either 0 if $i \neq j$ or the number of singletons person $i$ has otherwise. Together, all these facts about singleton $G$ mean the kinship is simply

$$K = N P_1^{\perp} diag(x) P_1^{\perp} \qquad (2)$$

where $diag(x)$ is a diagonal matrix with entries of $x$ on the diagonal.

Given this definition of K and the demeaned phenotypes $y_i' := y_i - \bar{y} = (P_1^{\perp} y)_i$ the H-E estimate is:

$$\hat{h}_{HE}^2 := \frac{Cov(K_{i>j}, y_i' y_j')}{Var(K_{i>j})}$$

where $Cov$ and $Var$ are the sample covariance and variance and the subscript on $K_{i>j}$ informally indicates the operators are applied to the (strict) lower triangle of the matrices $K$ and $y'y'^T$.

This definition is complicated and depends on everything but the variance terms given by the diagonal entries of $K$ and $y'y'^T$. But, intuitively, the diagonal terms of the inter-sample phenotypic and genetic similarity are all that matter—and this is formalized in Equation (1). Fortunately, H-E can still succeed because the demeaning operation spreads this diagonal self-similarity information across the entire $K$ and $y'y'^T$ matrices (e.g. Equation (2)).

Nonetheless, H-E regression seems unnecessarily complex, regressing on $\frac{N(N-1)}{2}$ data points when there are only $N$ independent observations. In fact, we show the H-E regression estimate is exactly proportional to the estimate from simply regressing $y'^2$ on $x$:

$$\hat{h}_{HE}^2 := \frac{1}{N-2} \frac{Cov(x, y'^2)}{Var(x)} \qquad (3)$$

This regression framework—which compares the singleton dosages to the size of demeaned phenotypes—is more intuitive than H-E, computationally simpler, more naturally models covariates, has analytic standard errors, can be generalized to appropriately weight the observations (reflecting the clear heteroscedasticity caused by singletons), and is easy to study theoretically.

The derivation details are below. The key fact is that both $K$ and $y'y'^T$ are demeaned versions of simple matrices—for $K$, it is a diagonal matrix, and for $y'y'^T$ it is a rank one matrix—but the calculations are tedious.

##### 2.4.2 Proving H-E is the same as regressing singleton count on squared phenotype.

First, the total off-diagonal kinship is

$$2\sum_{i>j} K_{ij} = \sum_{i,j} K_{ij} - \sum_{i} K_{ii} \quad = 0 - \; tr\big(N\, P_1^{\perp}\, diag(x)\big) = \; -N\, \frac{N-1}{N} \sum_{i} x_i \; = \; -(N-1)$$

Next, the total off-diagonal squared-kinship is

$$\frac{1}{N^2}\sum_{i,j} K_{ij}^2 = t\square(diag(x)\; P_1^{\perp} diag(x) P_1^{\perp}) = \sum_{i} x_i\, (P_1^{\perp} diag(x) P_1^{\perp})_{ii} \;\; \Rightarrow$$

$$\frac{2}{N^2}\sum_{i>j} K_{ij}^2 = \frac{1}{N^2}\sum_{i,j} K_{ij}^2 - \frac{1}{N^2}\sum_{i} K_{ii}^2 \quad = \sum_{i} x_i \left(\frac{N-2}{N}\, x_i + \frac{1}{N^2}\right) - \sum_{i}\left(\frac{N-2}{N}\, x_i + \frac{1}{N^2}\right)^2$$

$$= \sum_{i}\left(\frac{N-2}{N}\, x_i + \frac{1}{N^2}\right)\left(\frac{2}{N}\, x_i - \frac{1}{N^2}\right) \quad = \frac{N-2}{N}\frac{2}{N}\sum_{i} x_i^2 + \sum_{i}\left(-\frac{N-2}{N^3} + \frac{2}{N^3}\right) x_i - \frac{1}{N^3}$$

$$= \frac{2(N-2)}{N^2}\sum_{i} x_i^2 - \frac{N-3}{N^3}$$

We now combine these to get the variance of the lower triangle of $K$:

$$Var\big(K_{i>j}\big) = \frac{2}{N(N-1)}\sum_{i>j} K_{ij}^2 \; - \left(\frac{2}{N(N-1)}\sum_{i>j} K_{ij}\right)^2$$

$$= \frac{1}{(N-1)}\left(\frac{2(N-2)}{N}\sum_{i} x_i^2 - \frac{N-3}{N^3}\right) - \left(\frac{1}{N}\right)^2 \qquad = \frac{2(N-2)}{(N-1)}\, Var(x)$$

Now we need to compute to covariance term. First,

$$2\sum_{i>j} y'_i y'_j = \sum_{i,j} y'_i y'_j - \sum_{i} y_i'^2 = -\sum_{i} y_i'^2 = -N\, Var(y)$$

The inner product is

$$2\sum_{i>j} y_i' y_j'\, K_{ij} = \sum_{i,j} y_i' y_j' K_{ij} - \sum_{i} y_i'^2 K_{ii} = N\, y'^T diag(x) y' - \sum_{i} y_i'^2\left((N-2)x_i + \frac{1}{N}\right) = \sum_{i} y_i'^2\left(2x_i - \frac{1}{N}\right)$$

Together, this gives the covariance

$$\square ov\big(K_{i>j}, y_i' y_j'\big) = \frac{2}{N(N-1)}\sum_{i>j} K_{ij} y_i' y_j' \; - \left(\frac{2}{N(N-1)}\sum_{i>j} K_{ij}\right)\left(\frac{2}{N(N-1)}\sum_{i>j} y_i' y_j'\right)$$

$$= \frac{1}{N(N-1)}\sum_{i} y_i'^2\left(2x_i - \frac{1}{N}\right) - \left(\frac{-1}{N}\right)\left(\frac{-Var(y)}{(N-1)}\right)$$

$$= \frac{1}{N(N-1)}\sum_{i} y_i'^2\left(2x_i - \frac{2}{N}\right) \qquad = \frac{2}{N-1}\, Cov(y'^2, x)$$

Now Equation (3) follows by diving this by $Var(K_{i>j})$.

##### 2.4.3 H-E for Singleton LMMs is broadly unbiased

###### *2.4.3.1 Standard case*

Based on the equivalence to standard regression derived above, it is easy to prove H-E is approximately unbiased:

$$E\left( \hat{h}^2_{HE} \right) = \frac{1}{N} \frac{Cov(\, x, E(y'^2|x))}{Var(x)} = \frac{1}{N} \frac{Cov\left(\, x, x\, \sigma_g^2 + \sigma_e^2 \right)}{Var(x)} = \frac{\sigma_g^2}{N} \frac{Var(x)}{Var(x)} = \frac{\sigma_g^2}{N} = h^2$$

Again, informally, the $Cov$ and $Var$ operators integrate over the empirical distribution of $x$ while the $E$ operator integrates over random noise and singleton effect sizes. This assumes that the phenotypes $y$ have variance 1, which simplifies notation and makes the denominator in the definition of $h^2$ non-random, simplifying the expectation operations. More generally, the H-E estimate remains an unbiased estimate of the singleton variance explained.

###### *2.4.3.2 Random causals*

Imagine we observe $\dot{x}$ with $E(x|\dot{x}) = f\dot{x}$ for all $\dot{x}$; for example, only a fraction $f$ of the observed SNPs $\dot{x}$ are causal. Then

$$E\left( \hat{h}^2_{HE} \right) = \frac{1}{N} \frac{Cov(\, \dot{x}, E(y'^2|x))}{Var(\dot{x})} = \frac{1}{N} \frac{Cov\left(\, \dot{x}, E(x)\, \sigma_g^2 \right)}{Var(\dot{x})} = \frac{\sigma_g^2}{N} \frac{Var(f\dot{x})}{Var(\dot{x})} = \frac{f^2\sigma_g^2}{N}$$

While the genetic variance is attenuated from $\sigma_g^2$, this just reflects the fact that the true $x$ is $f$ times smaller than $\dot{x}$. In other words, H-E is still unbiased because

$$h^2 := E\big(Var(u|\dot{x})\big) + Var\big(E(u|x)\big) = E\big(x\, \sigma_g^2|\dot{x}\big) = \frac{f^2\sigma_g^2}{N}$$

###### *2.4.3.3 Stratified singletons: theoretical basis for partitioning singletons by global allele frequency*

In the analysis of our data, we partition singletons into multiple groups based on their global allele frequency (**Figure 2B/2C**, main text). We show in this section that this approach is well founded. In particular, we justify our approach to stratifying singleton heritability based on global allele frequency (as a proxy for the strength of selection acting on each causal singleton).

We generalize the standard LMM to have *K* distinct SNP groups, meaning that for each SNP $j$ in group $k$:

$\beta_j \sim N(\, 0, \frac{1}{M}\sigma^2_{g,k}\, )$ independently for all SNPs

The genetic contributions $u$ are still independent, but now have more complex variances. If $k(j)$ is the group for SNP $j$,

$$u_i := (G\beta)_i = \sum_{j=1}^{M} G_{ij}\beta_j \triangleq \sum_{j:G_{ij}=1} N\left(0, \frac{1}{M}\sigma^2_{g,k(j)}\right) \sim N\left(\, 0, \sum_{k=1}^{K} x_{ik}\sigma^2_{g,k}\right)$$

where $x_{ik}$ is the fraction of singletons that are both in group $k$ and carried by individual $i$.

Unfortunately, regressing $y'^2$ on $x$—i.e. H-E regression—no longer generally gives unbiased results. Note that the above random-causal SNP setting (2.4.3.2) is a special case of effect stratification with $K = 2$, but it is only because of the assumption that causals are uniformly distributed across observed SNPs that H-E is unbiased.

However, slightly generalizing H-E by regressing jointly on all singleton fractions—i.e. $X := (x_{,1}| \dots |x_{,K})$—gives unbiased estimates of the variance components:

$$E(y'^2|X) = \sum_{k=1}^{K} x_k \sigma_{g,k}^2 = x\,(\sigma_{g,1}^2| \dots |\sigma_{g,K}^2) \Rightarrow E(\,(X^T X)^{-1} X^T y'^2|X) = (\sigma_{g,1}^2| \dots |\sigma_{g,K}^2)$$

Of course, this requires knowing the true partition of the SNPs into the $K$ groups (or, slightly more generally, requires that the known partition of SNPs is finer than the true partition).

Again, if we condition on $Var(y) = 1$, the estimate of $h^2$ constructed from these unbiased variance component estimates is unbiased; more generally, it is asymptotically consistent and approximately unbiased.

###### 2.4.4 H-E is biased when the mean singleton effect is nonzero

We now modify the distribution on the genetic effects to have nonzero mean:

$\beta_j \sim N(\frac{1}{M}\mu, \frac{1}{M}\sigma_g^2)$ independently for all SNPs $j$

The genetic contributions $u$ are still independent with the same variances as before, but now they have nonzero means:

$$u_i := (G\,\beta)_i = \sum_{j=1}^{M} G_{ij}\beta_j \triangleq \sum_{j:G_{ij}=1} N\left(\frac{1}{M}\mu, \frac{1}{M}\sigma_g^2\right) \sim N(\,x_i\,\mu, x_i\,\sigma_g^2\,)$$

Under this non-standard LMM, it is easy to evaluate the expectation of the H-E estimate using the regression equivalence derived above:

$$NE(\hat{h}_{HE}^2) = \frac{Cov(\,x, E(y'^2|x))}{Var(x)} = \frac{Cov(\,x, V(y|x) + (E(y'|x))^2)}{Var(x)} = \frac{Cov(\,x, x\,\sigma_g^2 + \sigma_e^2 + \,(\mu x)^2)}{Var(x)}$$
$$= \sigma_g^2 + \mu^2 \frac{Cov(\,x, x^2)}{Var(x)}$$

The true heritability also changes from the addition of $\mu^2$:

$$h^2 = \frac{E(Var(u|x)) + Var(E(u|x))}{E(Var(y|x)) + Var(E(y|x))} = \frac{E(x\,\sigma_g^2) + Var(x\,\mu)}{E(x\,\sigma_g^2 + \sigma_e^2) + Var(x\,\mu)} = \frac{\sigma_g^2/N + \mu^2 Var(x)}{\sigma_g^2/N + \mu^2 Var(x) + \sigma_e^2}$$

Assuming the phenotypes have been standardized to variance 1, the denominator drops out, and the H-E bias due to mean singleton effects is

$$E(\,\hat{h}_{HE}^2) - \,h^2 = \frac{\sigma_g^2}{N} + \frac{\mu^2}{N}\frac{Cov(x,x^2)}{Var(x)} - \left(\frac{\sigma_g^2}{N} + \mu^2 Var(x)\right) = \mu^2\left(\frac{1}{N}\frac{Cov(x,x^2)}{Var(x)} - Var(x)\right)$$

When $\mu = 0$, this recovers our previous result that H-E is unbiased under the standard LMM. Otherwise, the bias scales with $\mu^2$ and a term comparing the second and third moments of $x$. This bias is negligible, though, as both terms in the difference are $O(x^2) = O\left(\frac{1}{N^2}\right)$.

**Table S2.** Simulation results showing that our new estimator of $h^2_{\text{singletons}}$ (SingHer $h^2$) is approximately unbiased across three distributions of the effect size (β), while in two scenarios H-E is biased. True $h^2$=0.3.

| β | SingHer $h^2$ | St. Err. | H-E $h^2$ | St. Err. |
|---|---|---|---|---|
| **random** | 0.2862 | 0.0041 | 0.2794 | 0.0064 |
| **fixed** | 0.2857 | 0.0034 | 0.7753 | 0.0082 |
| **biased** | 0.2825 | 0.0040 | 0.4499 | 0.0087 |

We performed simulations with true $h^2$=0.3 under three effect size distributions (**Table S2**), where random indicates $\beta$ is drawn i.i.d. from a mean zero Gaussian, fixed indicates $\beta_j = \mu$ for all $j$ (equivalently, a zero variance Gaussian), and biased indicates $\beta$ is drawn i.i.d. from a non-mean zero Gaussian. H-E returns reasonable estimates of $h^2$ if and only if the mean effect size is 0. This is consistent with our other simulations shown in **Figure 1A** (main text). We develop a novel model for jointly estimating $h^2$ and mean effect sizes that (approximately) solves this bias in the next section.

##### 2.4.5 Singleton LMMs can identify mean effect sizes

We have proved that non-mean zero genetic effect sizes, $\mu$, can badly bias H-E heritability estimates from singletons. It is not obvious how this meshes with the extensive publication record demonstrating the robustness of H-E. Indeed, ordinary H-E performs well in standard genome-wide simulations with $\mu \neq 0$ and $\sigma_g^2 = 0$[12].

The key difference is in $x_i$, the total minor allele dosage of person $i$. In the ordinary genome-wide context, where many SNPs are studied, $x$ is essentially constant across samples. In this case, a nonzero $\mu$ contributes roughly $\mu\bar{x}$ to each sample, which has two implications. First, the $\mu$ effect is indistinguishable from other sources of mean-shift in the phenotype. Second, the ordinary phenotype de-meaning step (in REML and H-E) appropriately residualizes $\mu$, and thus standard heritability approaches work well (in that the missed heritability, driven by $\mu$, will be negligible). However, in our setting—which involves a few (high-kurtosis) singletons in a *cis* window—$x$ has non-negligible variance: unfortunately, this biases standard heritability estimates; fortunately, it enables, for the first time, estimation of $\mu$.

We summarize our above non-mean zero singleton LMM as

$$y_i \sim N(\, x_i\, \mu,\; x_i\, \sigma_g^2 + \sigma_e^2\, )$$

and the $y_i$ are all independent.

We fit this model with REML, which projects out the intercept and $x$ from $y$. After this projection, we obtain the mean zero LMM again:

$$\breve{y}_i \sim N(\, 0,\; x_i\, \sigma_g^2 + \sigma_e^2\, )$$

where $\breve{y}$ are the residuals of $y$ after adjusting for the intercept and $x$; this is a common approximation and is correct to first order in $\frac{1}{N}$.

Interestingly, because the entries of $\breve{y}$ are independent, this likelihood problem is equivalent to the ordinary LMM likelihood after rotating away the kinship eigenvectors, with the singleton dosages $x_i$ replaced by the eigenvalues $\lambda_{\square}$ (note that eigenvalues can be interpreted as eigenvector dosages). In particular, any eigendecomposition-based LMM routine has a subroutine that fits the above likelihood model; we use the approach from[13]. We do not constrain the estimates of $\sigma_g^2$ and $\sigma_e^2$ to lie in (0,1), which enables approximately unbiased heritability estimates[14].

After fitting $\hat{\sigma}_g^2$ and $\hat{\sigma}_e^2$, we fit $\hat{\mu}$ by weighted least squares, which gives the ML estimate if $\sigma_g^2$ and $\sigma_e^2$ are estimated without error. Finally, we estimate the heritability by

$$\hat{h}^2 := \frac{\hat{\sigma}_g^2/N + (\hat{\mu}^2 + Var(\hat{\mu}))\; Var(x)}{\hat{\sigma}_g^2/N + (\hat{\mu}^2 + SE(\hat{\mu})^2)\; Var(x) + \hat{\sigma}_e^2}$$

This essentially evaluates the heritability definition at $(\hat{\mu}, \hat{\sigma}_g^2, \hat{\sigma}_e^2)$; the exception is the contribution of the standard error of $\hat{\mu}$, which reflects the fact that mean-zero error in $\hat{\mu}$ increases the expectation of $\mu^2$ beyond $\hat{\mu}^2$.

A different definition of heritability can be made removing the $Var(x)$ terms (from numerator and denominator). This is the ordinary REML definition, which is natural when the fixed effects are non-genetic[15]. However, in our model the mean effect is clearly attributable to genetic variation.

We note that because $E(y_i^2) = x_i^2 \mu^2 + x_i \sigma_g^2 + \sigma_e^2$, jointly regressing $y^2$ on $x^2$ and $x$ will yield unbiased estimates for $\mu^2$ and the variance components. We do not pursue this, however, because $\mu^2 < 0$ seems nonsensical; the computational cost of REML is negligible; and inverting an estimate of $\mu^2$ to estimate $\mu$ requires identifying the sign of $\mu$ and partitioning $\widehat{\mu^2}$ into $\hat{\mu}^2$ and $Var(\hat{\mu})$.

##### 2.5 Software availability.

Three open source software tools are being made available as part of this study, all available on GitHub:

HEh2.R – R code that performs all H-E analyses and simulations discussed in this paper. Also implements AI algorithm for parameter inference of LMM. Available here: https://github.com/hernrya/HEh2. Contact: Ryan Hernandez <>.

SingHer R package – Singleton Heritability inference with REML, discussed in Section 2.4, with performance statistics in **Table S2**, and available here: https://github.com/andywdahl/SingHer. Contact: Andrew Dahl <>, Noah Zaitlen <>.

Rejection sampling: Scripts demonstrating how we used rejection sampling to infer parameters of the phenotype model are available here https://github.com/uricchio/HE_scripts. Contact: Lawrence Uricchio <>.

### 3 GEUVADIS data set and QC

GEUVADIS: Genetic European Variation in Health and Disease, A European Medical Sequencing Consortium[16]. RNA-sequencing gene expression data were downloaded from www.geuvadis.org. This dataset contains 375 individuals of European descent from four locations. Each of these individuals are contained in the 1000 Genomes Project, and genome sequence data were downloaded from www.1000genomes.org[9].

##### 3.1 RNA-seq processing.

The GEUVADIS data consists of RNA-seq data for 464 lymphoblastoid cell line (LCL) samples from five populations in the 1000 genomes project. Of these, 375 are of European ancestry (CEU, FIN, GBR, TSI) and 89 are of African ancestry (YRI). In these analyses, we considered only the European ancestry samples. Some individuals were previously identified as having cryptic relatedness by the 1000 Genomes Project[9] using Identity by State (IBS) analyses and were therefore pruned. Our resulting dataset contains 360 unrelated individuals of European descent from four populations. Raw RNA-sequencing reads obtained from the European Nucleotide Archive were aligned to the transcriptome using UCSC annotations matching hg19 coordinates. RSEM [RNA-Seq by Expectation-Maximization, RSEM[17]] was used to estimate the abundances of each annotated isoform and total gene abundance is calculated as the sum of all isoform abundances normalized to one million total counts or transcripts per million (TPM). For each population, TPMs were log2 transformed and median normalized to account for differences in sequencing depth in each sample. The genotype data was obtained from 1000 Genomes Project Phase 3 V5 data set[9] . To remove potential confounders such as population structure and batch effects, we performed principal component analysis (PCA).

##### 3.2 PCA and Permutations: controlling for population structure and batch effects.

PCA was performed on both genome-wide genotype data as well as transcriptome-wide expression data. We obtained expression PCs from www.geuvadis.org, and ran PCA on the WGS data as follows. Input files:

1000 Genomes Phase 3 V5 variant call files

VCFtools [v0.1.14][18] to filter out related individuals, exclude singletons sites, remove indels, filter out all non-biallelic sites.

PLINK [v1.90b3x][19] was used to identify sites approximately in linkage equilibrium r^2 < 0.2 examining 50 kb windows in 5 site increments, extract these sites, and recode in an additive model (0, 1, 2).

R (https://www.r-project.org/) was used to concatenate chromosomes, and run principal component analysis on the centered and scaled genotype matrix.

We also ran PCA on the genotype data with a higher MAF filter (MAF≥5%) and got highly correlated results. However, because our analysis is based on rare variants, we wanted to include signals of population structure that manifest primarily in rare variants, hence including all variants seen at least twice.

We then checked for residual population structure using permutations. We first applied the standard permutation style, whereby phenotypes are shuffled among individuals prior to running H-E. We found that this unrealistically removed all signals in the data and gives $\widehat{h^2} = 0$. We then developed another permutation, which we refer to as a *trans*-permutation. In this case, we maintain the order of gene expression and genotypes among individuals, but we perform H-E regression on the SNPs in a window around one gene with the expression values of a random autosomal gene (i.e., a gene in *trans*). We show the results of this permutation in **Figure 1A** (main text) and in several figures below. We find that there exists some degree of residual population structure for rare variants, but not common variants (despite the fact that we included rare variants in our PCA analysis). The main caveat with this approach is that we are unable to distinguish population structure from rampant true *trans*-effects, but we feel removing the residual $\widehat{h^2}$ from the trans-permutation is conservative.

##### 3.3 Identifying high quality genes.

Not all genes are expressed in lymphoblastoid cell lines. To identify actively expressed genes, we analyzed several aspects of the data. **Figure S8** shows the distribution of $\log_2$(FPKM) across all individuals and genes. We then quantified the number of genes for which $X$% of individuals have $\log_2$(FPKM)>$Y$. For our main analysis, we focus on $X$=0.5 and $Y$=1.5, which results in a total 10,078, which we refer to as our Final Gene Set. **Figure S9** shows the cumulative proportion of $h^2$ as a function of MAF for various choices for $X$ and $Y$ (with $X$=0.5, $Y$=1.5 indicated in yellow). Each panel shows the number of genes

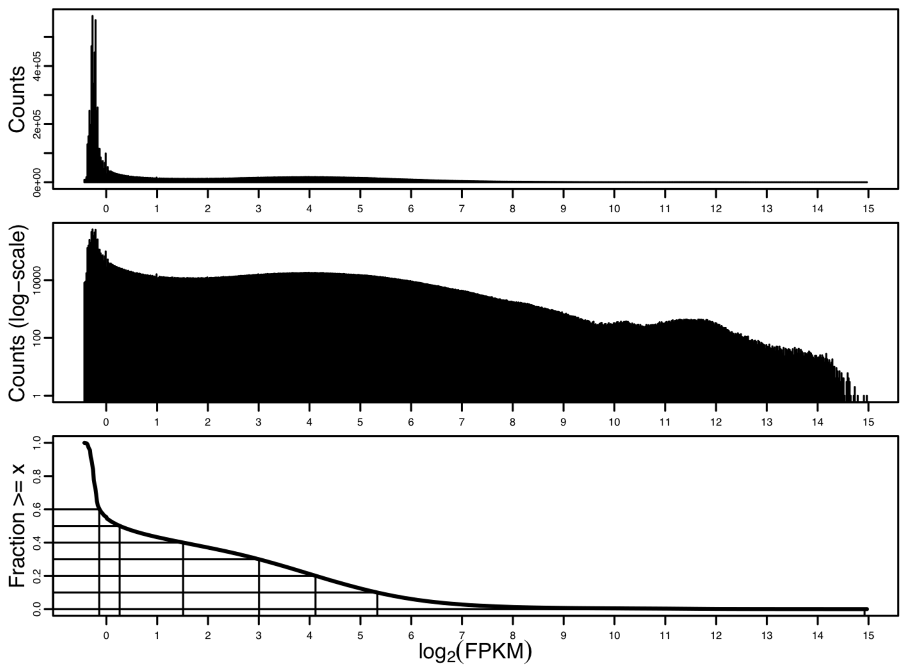


**Figure S8.** Investigating patterns of expression. Top shows the full distribution of all genes in all individuals. Middle is the same as t op, with y-axis on log scale. Bottom shows the inverse CDF, indicating how many observations (individuals×genes) have expression larger than the value on the x-axis. For reference 50% of all observations have $\log_2$(FPKM)≥0.26.

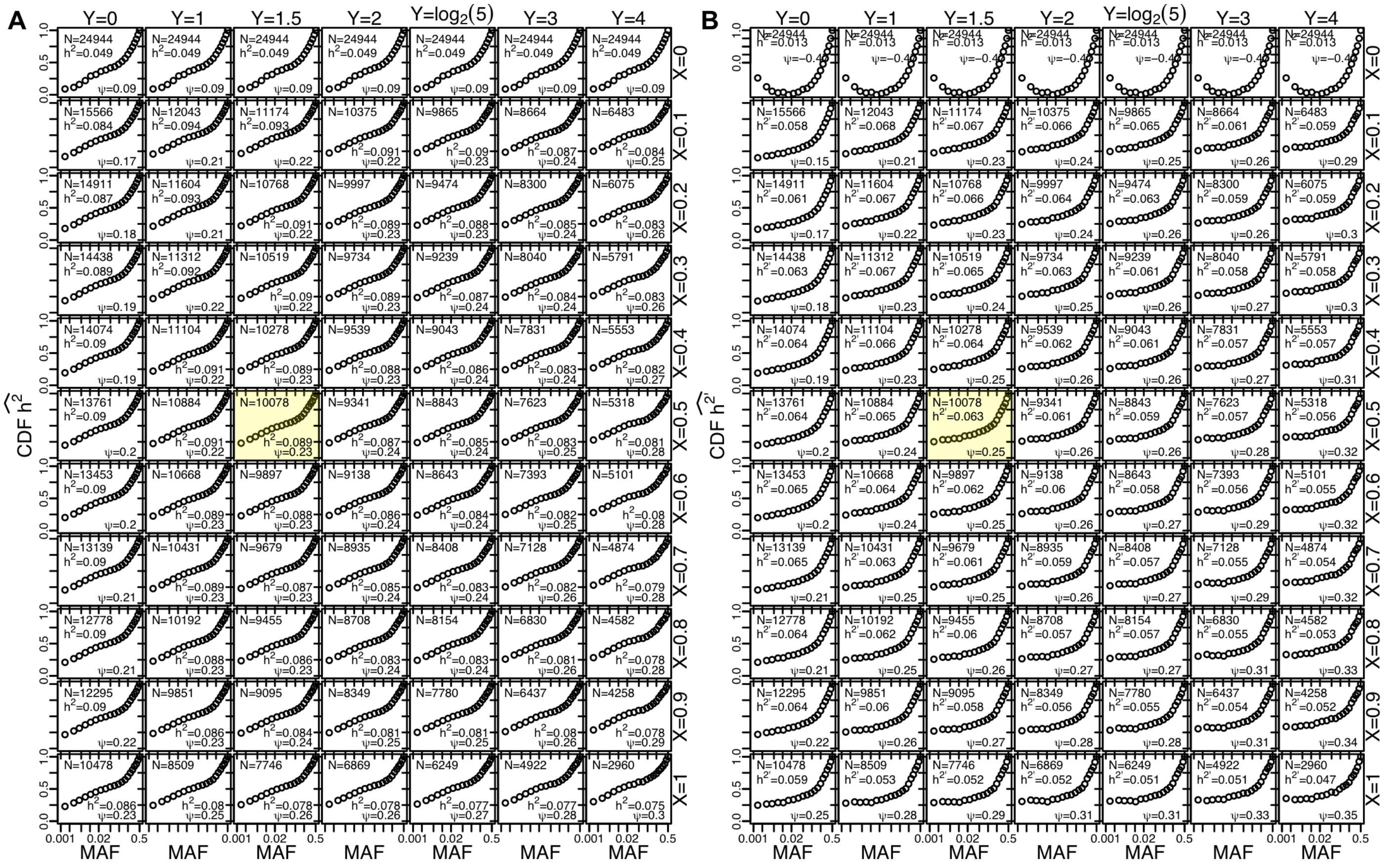


**Figure S9**. The cumulative proportion of heritability as a function of MAF for genes in which some fraction *X* of individuals (rows) have log2(FPKM)>*Y* (columns) with only PCA correction for population structure (**A**) and after correcting for population structure using our trans-permutation approach (**B**). For our main analysis, we focus on *X*=0.5 and *Y*=1.5, shaded yellow, but qualitative patterns are similar across all subsets except X=0, in which case heritability estimates for rare and low frequency variants are dominated by population structure. In each panel, N is the number of **autosomal** genes, $h^2$ is the total heritability estimate, and ψ is the fraction of heritability due to singletons.

that pass the X/Y threshold (N), the total heritability inferred ($h^2$), and the fraction of heritability explained by singletons (ψ). In general, these parameters do not impact our qualitative conclusions about the genetic architecture of gene expression so long as *X*>0. When *X*=0, many genes are effectively not expressed by any individuals, and both the estimate of $h^2$ and the fraction of $h^2$ due to singletons (ψ) are considerably lower (or negative). This figure also gives insight into the biological factors driving the genetic architecture of gene expression. Excluding the *Y*=0 column, we see that total $h^2$ monotonically decreases from left to right and top to bottom while ψ monotonically increases in both directions. This means that conditioning on some fraction of individuals having high expression reduces the total estimate of $h^2$, but also increases the impact rare variants have on patterning gene expression.

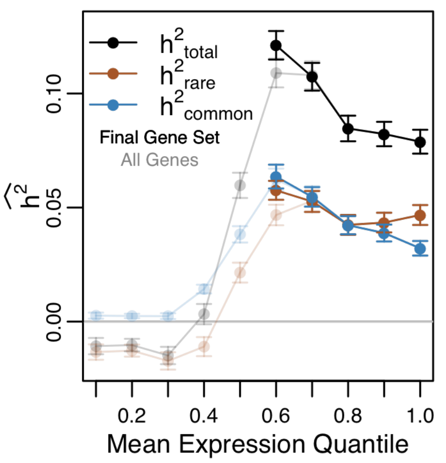


**Figure S10**. Mean (±2SE) total heritability inferred for genes with different expression levels (black) as well as the $h^2$ contributed by rare (MAF<=5%; red) and common (MAF>5%; blue) alleles. All genes were partitioned into disjoint decile bins, and the mean $h^2$ (total, rare, and common) for all genes in each bin is shown in lighter shades. Genes that make it into the final analysis are only in the upper quantiles, and shown in bold shades.

We further inspected the properties of $h^2$ inferred as a function of average expression levels across individuals for all genes, as well as those that make it in into our Final Gene Set. Specifically, we partitioned genes into decile bins based on their mean expression across all individuals (from lowest to highest expressed genes). In **Figure S10**, we show that for all genes (lighter shades), $h^2_{total}$ is near (or below) zero for the bottom forty percent of genes. Genes near the fiftieth expression percentile have positive $h^2$, but none of these genes made it into our Final Gene Set. All of the genes that made it into our Final Gene Set are in the 60th percentile and above. In general, we see that $h^2_{total}$ is maximized for genes with intermediate expression, where rare and common variants contribute in equal proportions to $h^2$. As we move toward higher expression quantiles, $h^2_{total}$ decreases.

##### 3.4 Robustness to model parameters.

In our inference of $h^2$, we make several model assumptions that could influence our inference of the genetic architecture of human gene expression. In the following subsections we present inference of $h^2$ as a function of MAF when we vary each of the assumptions made in our model, and argue that the assumptions we have made are in general conservative, and do not impact our qualitative conclusions.

###### 3.4.1 Number of SNP bins and Quantile Normalization.

Our primary analysis focuses on including all variants (down to singletons), K=20 SNP bins, and performing quantile normalization on the individual phenotypes. In **Figure 2** (main text), we show the dependence of our inference of $h^2$ on the MAF threshold. Adjusting the number of bins (from K=1 to 50) primarily affects our ability to make precise inference about the contribution of rare variants, since some rare variant classes tend to be pooled as K decreases (**Figure S11**). Subtle difference can arise (e.g., analyzing all SNPs in a single bin seems to result in a larger estimate of $h^2$, but, while K=2 results in a comparable $h^2_{total}$, the contribution of the MAF<1% bin seems to be underestimated). Quantile normalization plays a minimal role in our data analysis. This is in contrast to our simulations above (**Figure S7**), which suggest that when true $h^2$ is very large, quantile normalization can result in a substantial underestimate of $h^2$.

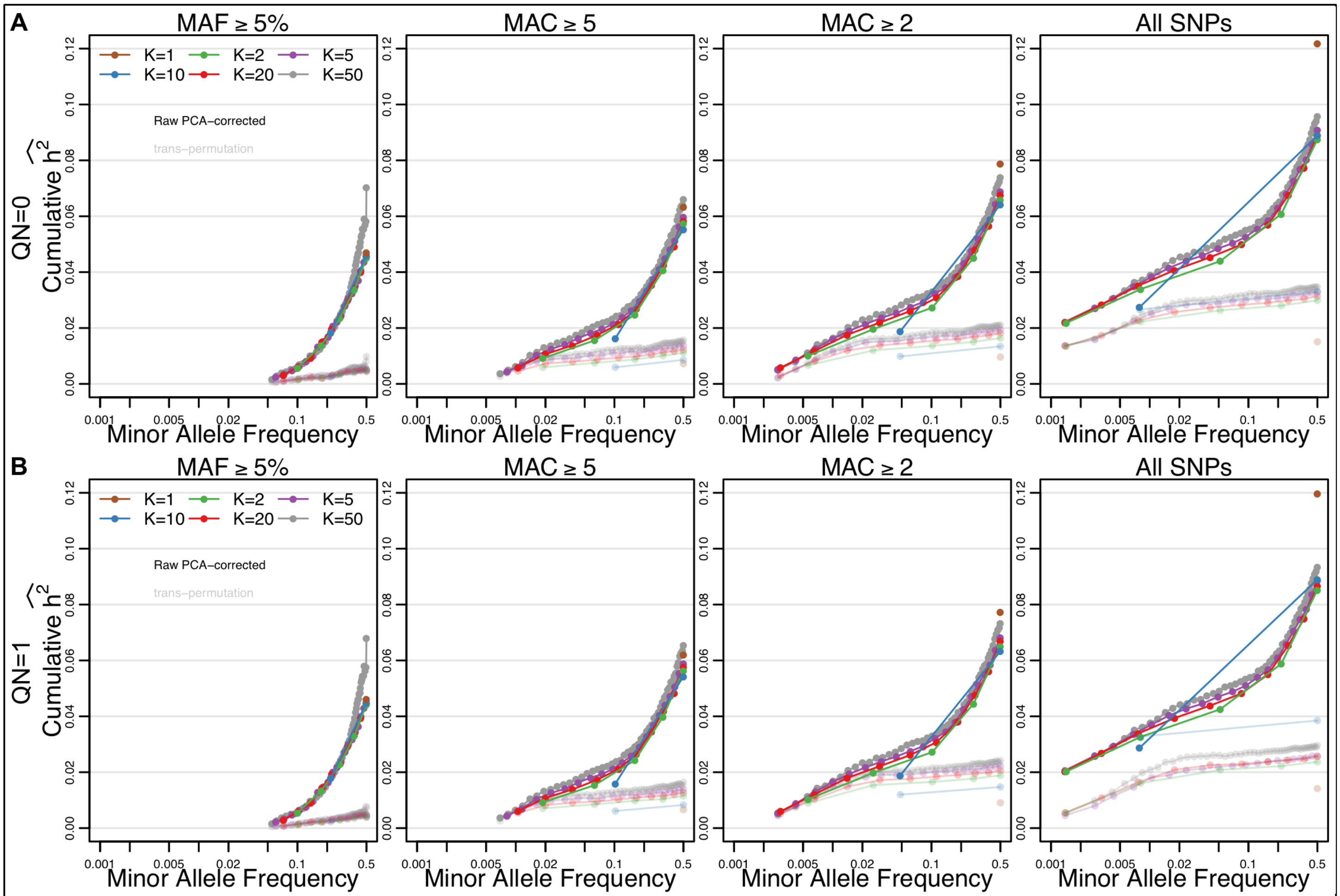


**Figure S11.** Effect of varying K and minMAF threshold without quantile normalizing the expression values (A) and with quantile normalizing expression values (B). In dark colors are the Raw PCA-corrected $h^2$ estimates, and in light shades are the estimates from the trans-permutation reflecting the degree of population structure.

##### 3.4.2 Window Size.

Our primary analysis focuses on a window size of 1Mb around the transcription start and end sites of each gene. We explored shorter windows with a length of 20kb and 100kb, and found comparable levels of $h^2_{\text{total}}$ (**Figure S12**). However, we enforce the constraint that each SNP bin must have at least 100 SNPs per gene to perform inference (otherwise adjacent bins are pooled together). As a result, when the window size drops from 1Mb to 100kb, we must drop to K=10. When the window size drops to 20kb, we must drop to K=5.

At shorter window lengths, the genetic architecture of gene expression is more dominated by common variation. This likely represents the substantial impact that nearby expression quantitative trait loci (eQTLs) play in shaping expression patterns among individuals (which are predominately common variants). Indeed, if we look at the pattern of $h^2$ as a function of MAF for genes that have a significant eQTLs in GEUVADIS data[20], we see that there is indeed a substantially larger contribution of common variants to $h^2_{\text{total}}$ (**Figure S13**). Interestingly, while common variants play a larger role in genes with an eQTL, rare variants contribute the same level of $h^2$ across both eQTL and non-eQTL genes. As the eQTLs were identified in this data, there could be a winner's curse that results in an overestimation of $h^2$ for common variants, and not necessarily predictive of the patterns in independent datasets.

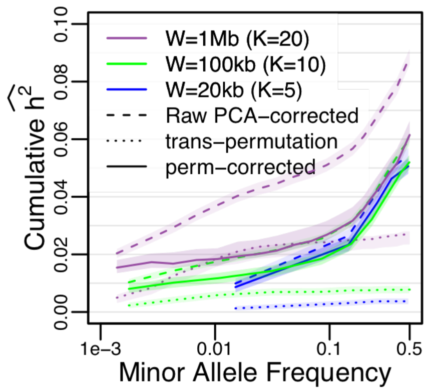


**Figure S12.** Effect of window size on inference of genetic architecture. Shorter windows around each gene results in smaller numbers of SNPs, which reduces the K we can use, but has only a moderate impact on $h^2_{\text{total}}$. Rare variants play a large role outside the immediate vicinity of genes. 95% bootstrap confidence envelopes are shown.

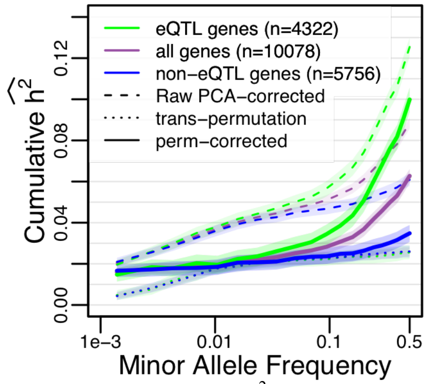


**Figure S13.** Cumulative $h^2$ as a function of MAF for all genes versus those genes that have or do not have a genome-wide significantly associated eQTL. 95% bootstrap confidence envelopes are shown.

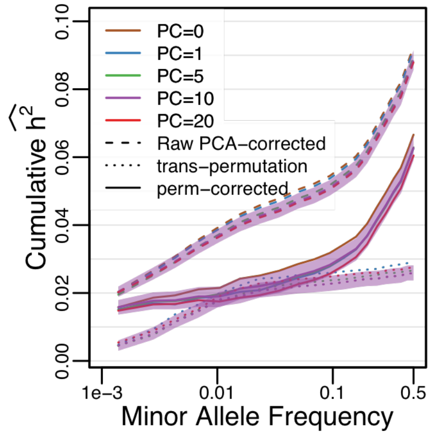


**Figure S14.** Adjusting the number of principal components does not affect our inference of $h^2$ or the contribution of rare variants. 95% bootstrap confidence envelopes are shown.

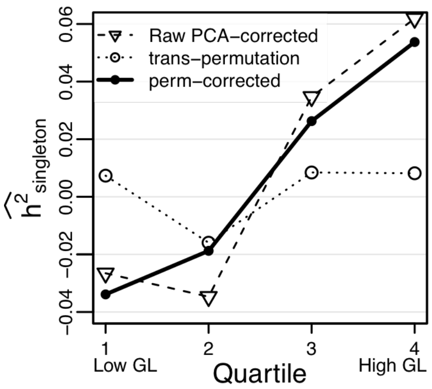


**Figure S15.** Partitioning singletons in our data into quartile bins based on genotype likelihood demonstrates that not all singletons contribute equally to the inferred heritability. Singletons with higher genotype likelihood contribute more to inferred heritability than singletons with low genotype likelihood.

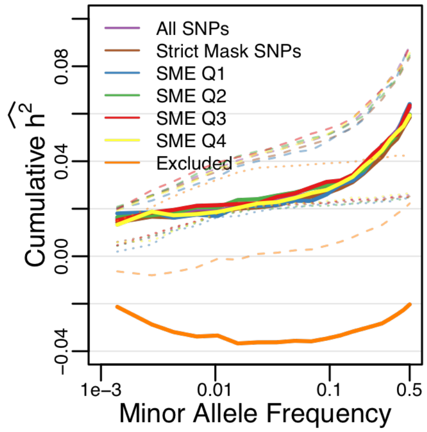


**Figure S16.** Partitioning genes by the proportion of TGP Strict Mask exon (SME) bases suggests that all genes included in our analysis have result in the same global pattern. In contrast, genes excluded from our analysis (orange) are substantially different.

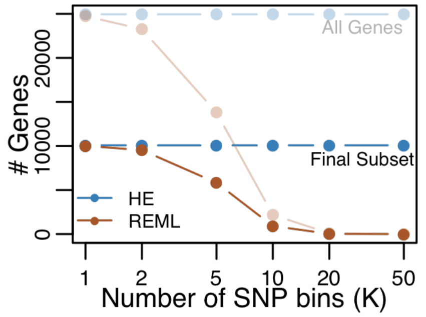


**Figure S17.** The number of genes that converged for H-E vs LMM (REML) for All Genes (light shades) or the Final Gene Set (dark).

##### 3.4.3 Number of PCs.

The bulk of our analysis focuses on the inclusion of the top 10 principal components from both genotypes as well as phenotypes. These are intended to eliminate possible confounding effects of population structure (in the genetic data) or batch effects (in the RNA-seq expression data). How PCs are added to the H-E regression model is discussed above. The number of principal components used makes very little difference to our inferred signature, though $h^2_{total}$ decreases very slightly as the number of PCs increases (**Figure S14**). Note that the residual population structure we identify with trans-permutations is not affected by PC correction.

#### 3.5 Genotype quality and genome mapability.

One potential concern is that regions of the genome could contribute to poor quality read mapping, and this could confound both genotype data (SNP calls from predominantly low coverage whole genome sequence data) as well as expression data (measuring FPKM with RNA-seq). To critically assess these issues, we performed two additional experiments. We first interrogated our SNP calls, particularly singletons since they are a major source of low quality genotype calls (and the largest source of $h^2$). For each singleton, we extracted the genotype likelihood of the individual who carried the minor allele from the 1000 Genomes Project (TGP) supporting information (ftp://ftp.1000genomes.ebi.ac.uk/vol1/ftp/release/20130502/supporting/genotype_likelihoods/). We then divided these singletons into four disjoint quartile groups based on their genotype likelihood (sorted from lowest quality to highest quality), and performed H-E regression on these four groups of singletons. In **Figure S15**, we show that singletons in the lowest genotype likelihood quartile contribute nothing (or slightly negatively) to $h^2_{singleton}$. Instead, only the upper quartiles contribute positively to $h^2_{singleton}$. This is in contrast to partitioning singletons by their global allele frequency observed in gnomAD, which shows that the lowest frequency bins contribute the most to $h^2_{singleton}$ (**Figure 2B**, main text). This is strong evidence that SNP genotype quality is not inflating our signal of $h^2$.

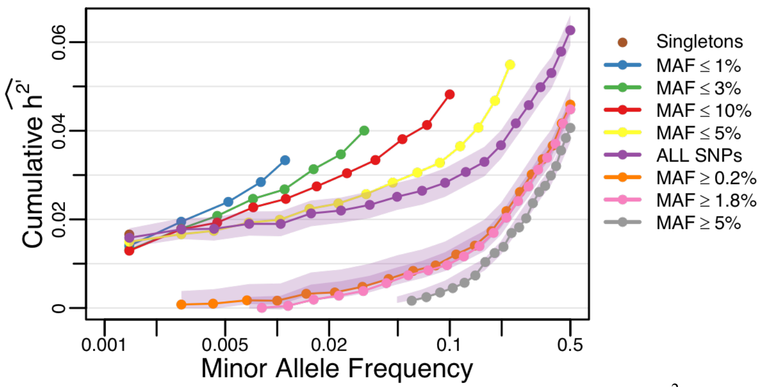


**Figure S18. Tagability of rare and common variants.** Cumulative $h^2$ inferred for multiple subsets of SNPs. In the middle are the purple dots, which includes all data (as shown in **Figure 2C**, main text). Above are cumulative $h^2$ estimates excluding common variants to ask whether rare variants tag common variants. That these curves tend to deviate above the purple dots indicates that rare variants can partially tag the $h^2$ of common variants. Below the purple dots are cumulative $h^2$ estimates excluding rare variants to ask whether common variants tag $h^2$ from rare variants. The purple shaded area indicates the 99% quantile range of the mean for 1,000 bootstrap samples of the full data set. From this envelope, we subtracted the contribution of rare variants, and find that it largely overlaps the inferred $h^2$ when rare variants are not included in the model. This demonstrates that common variants do not tag any rare variant $h^2$.

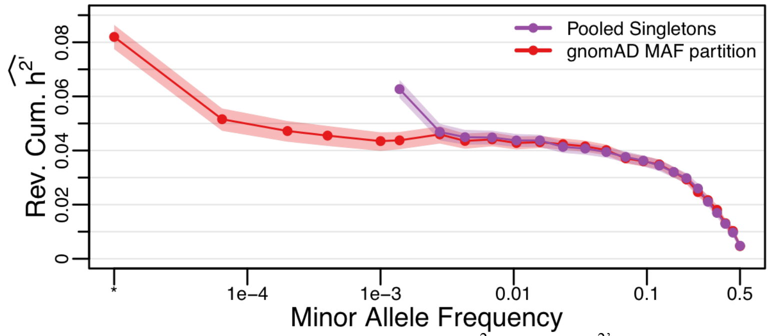


**Figure S19.** Reverse cumulative distribution of $h^2$ (i.e. total $h^{2'}$ contributed by variants above a given MAF) when singletons are pooled (purple) vs partitioned according to their observed gnomAD MAF (red). 95% bootstrap confidence envelopes are shown.

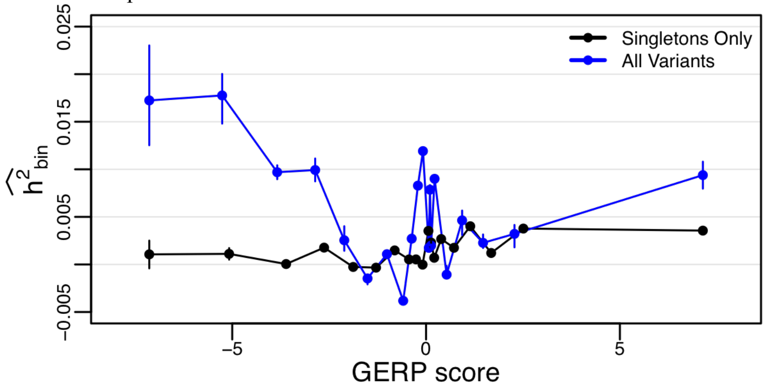


**Figure S20. Heritability for SNPs with different evolutionary constraint.** We partitioned SNPs into quantile bins by their evolutionary constraint as measured by GERP (where negative values indicate more cross-species constraint and positive values indicate rapid evolution). We partition singletons only (black), and all SNPs (blue) by GERP. This suggests that evolutionary constraint may be more helpful for common variants than for rare variants. 95% bootstrap confidence intervals are shown.

We next interrogated the possibility that difficult to sequence regions of the genome could be leading to read mapping errors for the RNA-seq data, shown in **Figure S16**. We first reproduced the main analysis of all 10k genes that are in our final analysis (purple curve, which you cannot see because it is completely covered by other curves). We then re-analyzed all genes, but included only those SNPs that were in regions of the genome passing the TGP accessible genome strict mask (ftp://ftp.1000genomes.ebi.ac.uk/vol1/ftp/release/20130502/supporting/accessible_genome_masks/), shown in brown in **Figure S16** (again, difficult to see because it is buried by other curves). We next partitioned genes into four quartile bins based on the number of exon bases that were contained in the strict mask (Q1 having the lowest fraction, Q4 having the highest fraction). The idea is that regions of the genome that are difficult to map could result in aberrant RNA-seq expression values. We find no evidence of this. The genetic architecture in each partition is essentially indistinguishable (**Figure S16**, blue, green, red, and yellow curves). This is in contrast to the genetic architecture of the genes that were excluded from our analysis due to insufficient expression across enough individuals [i.e., <50% of individuals had $\log_2$(FPKM)>1.5, shown in orange in **Figure S16**].

Together, these observations strongly suggest that data quality has not compromised our inference of the genetic architecture of human gene expression.

#### 3.6 Comparing H-E and REML

A majority of papers analyzing $h^2$ from unrelated individuals focus on linear mixed models (LMMs) using REstricted Maximum Likelihood (REML), and its extensions. We found that we are unable to obtain reliable inference of $h^2$ from REML due to lack of convergence at our sample size. This problem becomes particularly problematic when the number of SNP bins increases. Shown in **Figure S17** is the number of genes for which we are able to infer $h^2$ from H-E (blue) and REML (brown), for both the total set of all genes (light shades) as well as the Final Gene Set (dark shades). Essentially all genes fail to converge with K=20 despite multiple independent attempts for each gene.

#### 3.7 Common variants tag a negligible amount of $h^2$ from rare variants

The extent to which common variants are able to tag heritability contributed by rare variants (and vice versa) is a major open question in the field. To investigate this question, we analyzed multiple subsets of SNPs as a function of MAF (**Figure S18**). We start by reproducing the plot of cumulative $h^2$ as a function of MAF for the full data set (purple, **Figure S18**, as shown in main text **Figure 2C**). We then inferred cumulative $h^2$ as we removed progressively more common variants (curves above purple dots in **Figure S18**). We find that each of these curves fall above the purple dots, which demonstrates that low frequency variants are able to partially tag $h^2$ contributed by more common variants. We also performed the reverse exercise, where we strip out progressively more rare variants, which as shown in **Figure 2C** (main text) results in a reduced $h^2_{total}$ (below purple dots). To evaluate whether some of the rare variant $h^2$ was being tagged by common variants, we performed the following analysis. We calculated the cumulative $h^2$ as a function of MAF for 1,000 bootstrap samples of all genes in our final gene set. For each bootstrap sample, we removed progressively more rare variants, and plotted the 99% quantile range across bootstrap samples with a purple envelope. **Figure S18** shows that this purple envelope covers the orange, pink, and grey points, demonstrating that common variants have not tagged any $h^2$ contributed by rare variants.

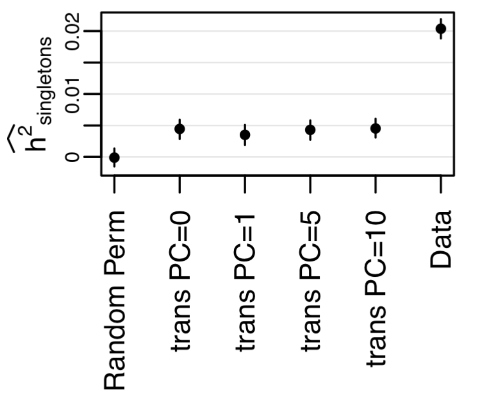


**Figure S21. Population structure is a negligible contributor to singleton heritability.** We ran permutations multiple ways to characterize the impact of population structure on singleton heritability. Points correspond to the mean across all genes ±95% quantile range of 10,000 bootstrap samples across genes. See Section 3.9 for description.

We further evaluated whether partitioning singletons by gnomAD MAF resulted in additional common variant $h^2$. To do this, we recreated the red and purple curves shown in **Figure 2C** (main text) using a reverse cumulative distribution (**Figure S19**), which shows the total $h^{2'}$ contributed by variants with $MAF \geq x$. This shows that common variant $h^2$ is unaffected by singleton partitioning, suggesting that common variant $h^2$ has not been impacted by this partitioning scheme.

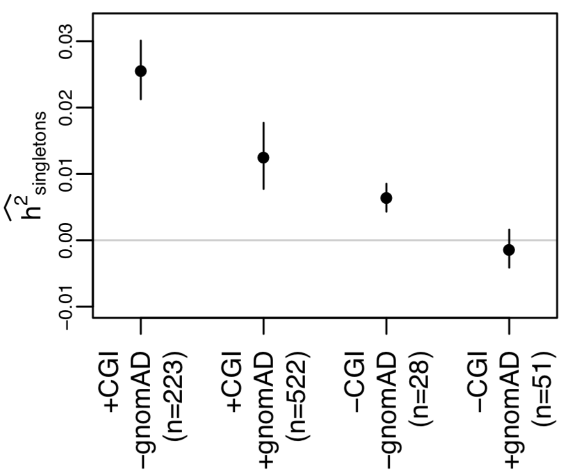


**Figure S22. Partitioning singletons by presence or absence in high coverage data and the gnomAD resource.** We find that the vast majority (88%) of our singleton $h^2$ reported in **Figure 2** (main text) are contributed by variants that are validated in high coverage sequencing data. Further, the majority of this signal (67%) is contributed by singletons that are not in gnomAD. 95% bootstrap confidence intervals are shown.

##### 3.8 Heritability as a function of evolutionary conservation.

We inferred an evolutionary model for the genetic architecture of gene expression, and it suggests that causal variation is constrained by the action of natural selection (**Figure 3**, main text). It is therefore plausible that evolutionary constraint as measured by comparative genomics using the GERP statistic[21] could aid in the identification of causal variation. In **Figure S20** we show that for singletons, sites with stronger evolutionary constraint (i.e. sites with more negative GERP scores) do not in general have increased $h^2$. In contrast, when all sites are partitioned by their GERP score, evolutionary constrained sites do contribute substantially more $h^2$. This suggests that evolutionary conservation may be an important metric for common variants, but not as informative for singletons. This could be because the regulatory architecture of the human genome has evolved sufficiently quickly that long-term constraint does not capture all aspects of functionality in modern humans.

##### 3.9 Population structure makes a negligible contribution to singleton heritability.

Previous work has shown that inclusion of PCs can remove population structure bias from common variant estimation. We included PCs in our inference of $h^2$ and here evaluate whether there is residual population structure contributing to our inference of $h^2_{singletons}$. We performed multiple permutations, and show in **Figure S21** that none of them produce substantial singleton heritability. First, a typical permutation would shuffle the genotype-phenotype relationship among individuals. Such a permutation results in zero $h^2$ estimates ("Random Perm" in **Figure S21**). However, population structure in our sample renders such permutations inadequate. Our data analysis utilizes PCA to control for population structure, but since genotype-phenotype relationships are permuted, PCA-correction can no longer be easily applied [though other approaches are possible[22]]. We therefore constructed an additional permutation, where we paired the genotypes around one gene with the expression values of a random gene from the same individual. Since the individuals are preserved in this permutation, PCA can be applied, but we find that it has no effect of $h^2_{singletons}$, which are all between 0.0035-0.005 ("trans PC={0,1,5,10}" in **Figure S21**). These estimates of singleton heritability from trans-permutations are negligible compared to the real data ($h^2_{singletons}$=0.02 for the GEUVADIS MAF partition, labeled "Data" in **Figure S21**), so we are therefore confident that population structure has a minimal impact on the patterns of $h^2_{singletons}$ that we infer from our data. Work from other groups has considered the possibility of fine-scale populations not captured by PC covariates. In this case our heritability estimates, as well as previous estimates[2,3], as well as tests for association to rare variants, may be slightly inflated.

##### 3.10 Analyzing a subset of individuals with high coverage whole genome sequencing

As part of the TGP[9] analysis, a subset of individuals were sequenced to high coverage by Complete Genomics Inc. (CGI). Fifty-eight of these subjects overlap the individuals in our study from GEUVADIS. To gain insight into the role of false-positive and false-negative inference of singletons in our estimation of $h^2$, we partitioned the singletons observed in the 1000 Genomes Project low coverage data by their presence/absence in both the high coverage CGI data as well as gnomAD. We find that the vast majority (88%) of the singleton heritability that we report in **Figure 2** (main text) are contributed by singletons that replicate in the high coverage data (**Figure S22**). Further, a majority of this signal (67%) is contributed by variants that are not present in the gnomAD dataset.

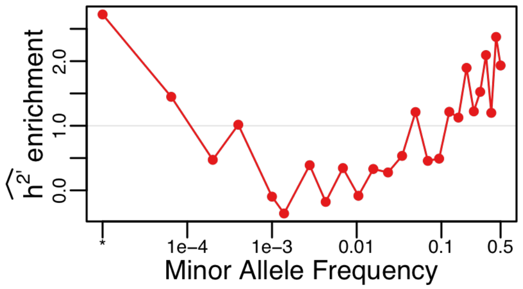


**Figure S23.** Enrichment of heritability across MAF bins, calculated as the average fraction of bias-corrected inferred h2 per bin across genes (as shown in main text Figure 2C) divided by the average fraction of variants observed in each MAF bin.

#### 3.11 Heritability enrichment across MAF.

In **Figure 2** (main text) and above, we show that rare variants (particularly singletons that are ultra-rare in gnomAD) compose a substantial fraction of heritability. It is also the case that the plurality of variants in *cis*-regions of genes are rare. Oppositely, common variants compose a majority of the remaining heritability, but, following population genetic theory, there are fewer common variants than rare variants in *cis*-regions of genes. We therefore performed an enrichment analysis to see if there is more heritability inferred in a MAF bin than expected from the fraction of variants observed across MAF bins. In **Figure S23,** we show that heritability enrichment is “U” shaped as a function of MAF (on a log scale). This suggests that the rarest and most common alleles contribute an excess of heritability, and intermediate/low frequency variants (MAF 0.1%-5%) compose a dearth of heritability. While this does not give us direct insight into the underlying distribution of regulatory effect sizes per causal variant, it may be reasonable to speculate that in order for the rarest alleles to have the largest enrichment, the distribution of effect sizes for rare causal variants must be considerably larger (in absolute value) than common variants. We explore this topic further below with model predictions in Section 4.

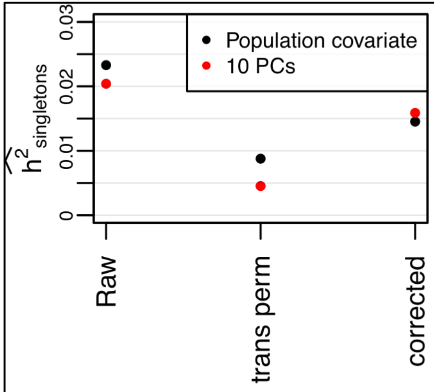


**Figure S24.** Comparing the use of a series of dichotomous population covariates to control population structure instead of PCA. Both approaches result in comparable levels of singleton heritability inferred after correcting with our *trans*-permutation.

#### 3.12 Exploring population covariates as an alternative to PCA.

In much of our analysis, we focus on PCA to control for population structure (as described in Section 3.2). In reality, we know that our samples derive from four different European populations, and therefore explored the use of a series of binary covariates to indicate the population of origin for each sample. **Figure S24** shows that both approaches produce comparable results. Though the population covariates resulted in slightly higher estimates of singleton heritability, this was compensated by slightly higher residual population structure from our *trans*-permutation test.

#### 3.13 Constructing bootstrap confidence intervals.

In main text **Figure 2** and throughout this Supplemental Material, we compare heritability estimates in many ways. Our primary approach to estimating uncertainty was based on rigorous bootstrapping. Except where noted, all error bars (sometimes plotted as envelopes encompassing the mean) were calculated from the 95% interquantile range of 1,000 bootstrap samples. This is an appropriate method for estimating uncertainty in i.i.d. data, and has previously been shown to work well in far broader settings[23]. Further, bootstrapping is a statistically appropriate way to estimate uncertainty when analyzing functions of correlated parameter estimates (for example, when estimating total $h^2$, which is the sum across $h^2$ estimates per bin). These bootstrap intervals represent uncertainty in the across-gene average heritability estimates per category (indeed, the single gene uncertainties are much larger), and retains any across-gene correlations that are present in the real data. Hence, our SE estimates naturally account for correlated expression.

### 4 Inferring posterior distributions of evolutionary parameters

#### 4.1 Overview of rejection sampling pipeline

We first provide a high-level overview of our rejection sampling[24] procedure in this section, and in the subsequent sections provide in-depth details pertaining to our model and its implementation.

Rejection sampling compares a set summary statistics that are informative about model parameters computed on the output of model-based simulations to summary statistics computed on observed genomic and phenotypic data. The simulations that generate summary statistics that are most similar to the observed data are retained, and the parameter values from the retained simulations are used to generate a posterior distribution over the true parameter values.

The goal of our rejection sampling procedure is to infer the mean strength of selection (*2Ns*, where *N* is the effective population size and *s* is the fitness effect; represented here by a parameter $\phi$ that corresponds to a mixture proportion of weakly and strongly selected sites), the correlation between selection on variants and their effect on fitness ($\rho$), and the mean shape of the effect size distribution ($\tau$), for the alleles that contribute to gene expression variation among individuals. Details on the model were previously described[5] and summarized below. We focus on inferring the posterior distribution of the mean across genes as opposed to the parameter values for individual genes because single gene estimates proved too noisy to be reliably computed.

Our estimation procedure has 4 main steps, which we then further subdivide into several small steps in the subsequent sections. The first main step is to generate 2,000,000 simulated gene expression values using parameter values ($\rho$, $\tau$, $\phi$) sampled from prior distributions. For our prior distributions, we use a Beta(0.35, 0.35) distribution for $\tau$ and $\rho$ and a Uniform(0,1) distribution for $\phi$, such that there are an excess of genes with very large and very small $\tau$ and $\rho$, and selection strengths are evenly distributed between 0 and 1 (note that the prior distributions over *gene sets* that we use for inference are uniform, while the prior over individual simulated genes were not uniform for $\tau$ and $\rho$). These simulated genes utilize forward-in-time simulations of selection and human demography to generate a distribution of selection coefficients and allele frequencies, then use an evolutionary phenotype model to map these selection coefficients to effect sizes, and finally apply these effect sizes to empirical *cis*-variation for each sequenced gene used in the data analysis to generate phenotypes for each individual (along with an error term representing random environmental effects).

Second, we perform H-E regression on each simulated phenotype with real genotypes, and compute cumulative $h^2$ as a function of MAF for each simulated gene (*i.e.*, the same analysis that we have performed for each gene in the main text for the observed GEUVADIS data).

Third, we resample 1,000,000 gene sets, each composed of 10,000 genes, from the complete set of 2,000,000 simulated expression products. We use 10,000 genes to approximately match the number of genes in the final gene set we analyze in the observed GEUVADIS data. The genes are resampled such that the mean $\tau$, $\rho$ and $\phi$ over the gene set are approximately equal to value of $\tau$, $\rho$ and $\phi$ drawn from a uniform prior distribution for each parameter (note again that the prior distributions over *gene sets* are uniform, while the prior over individual simulated genes were not uniform for $\tau$ and $\rho$).

Lastly, we compute mean cumulative $h^2$ as a function of MAF over each of the 1,000,000 simulated gene sets, and input this set of parameter values (mean $\tau$, $\rho$ and $\phi$) and paired summary statistics (cumulative value of $h^2$) into our rejection sampler, which computes the mean squared distance between the observed and simulated $h^2$ vs

MAF curve for each gene set. We then retain the 1,000 gene sets that minimize this distance to form an approximate posterior distribution.

Below, we provide a more detailed description of each step.

#### 4.2 Evolutionary model of complex phenotypes

In order to simulate phenotypes, we must map simulated selection coefficients to causal alleles with a given effect size. We employed a previously developed evolutionary model[5] of phenotypes under selection that captures the relationship between effect sizes and selection strength as

$$\beta = \begin{cases} \delta|s|^{\tau} & \text{with probability } \rho \\ \delta|s_r|^{\tau} & \text{otherwise} \end{cases}$$

where $s$ is the selection coefficient of a causal site while $s_r$ is a random selection coefficient drawn from the marginal distribution of selection coefficients over all causal sites; $\rho$ controls the correlation between the strength of selection on the variant and the phenotypic effect of the variant; $\delta$ is a random sign; and $\tau$ is an exponent that controls the shape of the relationship between selection coefficients and effect sizes.

Conceptually, if ρ is near 1, then the effect that alleles have on gene expression are tightly correlated with their selection coefficients, which would imply that strong effect mutations tend to be evolutionarily deleterious. If ρ is closer to 0, then for the majority of causal alleles, effect sizes are disjoint from the strength of selection. Such cases could reflect a high level of pleiotropy, whereby alleles with weak effects could be experiencing strong selection because of other functional effects of the allele (and vice versa). If τ is close to 0, then all effect sizes are of comparable magnitude, whereas if τ is large, effect sizes are broadly distributed and large effect alleles are not unlikely. Unlike ρ, τ is not formally constrained on [0,1], but in practice this range is suitable since the distribution of fitness effects that have been inferred for humans tends to be very widely distributed[25,26], and previous analyses inferring τ have found values between 0 and 1.[2] For more discussion of the model, see[5].

#### 4.3 Forward in time-simulations

Since population demography and natural selection both alter genetic architecture[5,27–29], we performed forward-in-time simulations under a population genetic model of recent human demography[30] and fitness effects[25,26] using SFS_CODE and sfs_coder[31,32]. The demographic model includes an expansion in the African ancestral population, an out-of-Africa bottleneck, a second bottleneck at the founding of Europe, and followed by super-exponential growth. Parameter values of the simulations are reported elsewhere[5,32].

To model natural selection, we supposed that alleles that alter the expression levels of genes would be composed of a mixture of strongly selected and weakly selected loci. We therefore simulated a mixture of alleles under strong selection [consistent with human nonsynonymous sites, $\langle 2Ns \rangle$= -457][26] and alleles under weaker purifying selection [consistent with conserved non-coding sites, $\langle 2Ns \rangle$ = -8][25]. Note that this mixture parameter is one of the parameters that we infer with our rejection sampling procedure, and hence our method allows for a wide range of selection strengths that are consistent with previous estimates of selection in humans. Moreover, since the selection and demographic parameters were inferred from patterns of diversity in human genomic sequences, our simulations will generate patterns of diversity that are broadly consistent with human patterns of polymorphism in functional regions.

At the end of each simulation, we down-sampled to 360 individuals to match the observed GEUVADIS data, and then used sfs_coder to extract the simulated allele frequencies and selection coefficients[32].

#### 4.4 Simulated phenotypes

Our simulated phenotypes use the actual sequence data from GEUVADIS by matching each observed allele to an effect size (many of which are 0). For each simulated gene, we first sample values of τ, ρ and ϕ from prior distributions as described above. For each set of parameter values, we then sample causal variants from the simulated frequency spectrum, and then map the selection coefficient of this allele to an effect size under our model. We then sample a frequency-matched allele in the observed genomic data using the parameters of our evolutionary phenotype model (discussed above). We assume that between 2% and 20% of the sites surround each gene are causal for the phenotype. We selected 2% and 20% to reflect the diversity of conserved non-coding sequence density in the human genome, supposing that functional sites are likely to have an effect on expression (although this effect may be very weak, depending on the parameters of selection model).

Once effect sizes have been sampled, we computed the genetic component of the phenotype under a standard additive phenotype model (*i.e.*, by summing up the effect sizes of each variant for each individual in the analysis). We then simulated a random normal variable to represent the environmental contribution to the variance in expression level for each gene. The random normal variable was scaled such that the true narrow sense heritability of each simulated expression level was set to $h^2$= 0.09, in concordance with the approximate mean heritability observed in our allele frequency partitioned analysis of GEUVADIS expression levels. Finally, we quantile normalized the simulated expression of each gene.

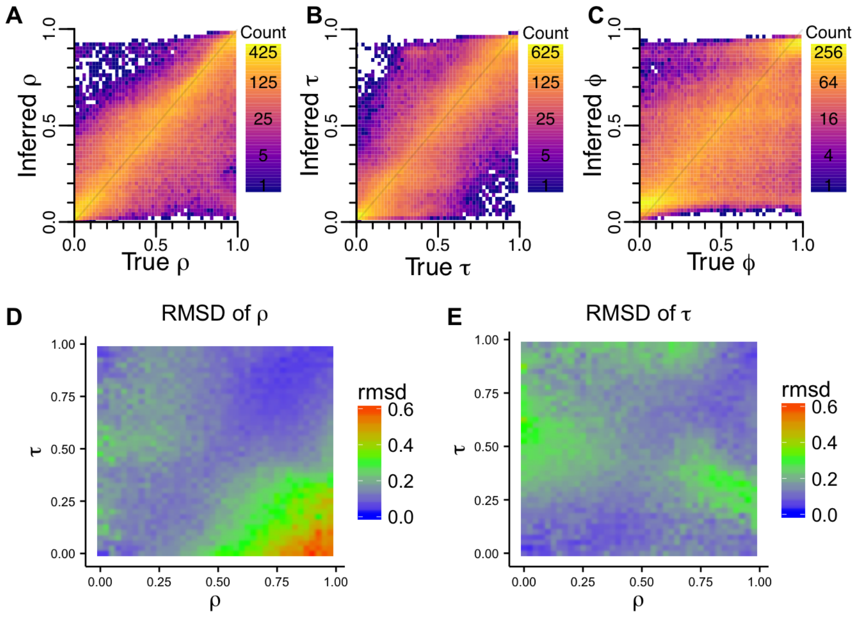


**Figure S25.** Model performance for rejection sampler. (A-C) shows the distribution of inferred vs the simulated parameters, demonstrating a strong correlation (note that the coloration is on a log scale, indicating that inference is tightly distributed along the diagonal). However, performance is non-uniform. Here we plot the root mean squared deviation (RMSD) between true and inferred values of ρ (D) and τ (E) across the parameter space as a function of the other parameter. RMSD tends to be minimized along the diagonal (e.g. τ≈ρ). When τ is very small, our model has difficulty inferring large ρ (note that τ=0 forces all causal variants to have the same effect size, so it does not make sense to have a correlation between fitness effects and phenotype effect size if all causals have the same effect size).

#### 4.5 H-E on simulated expression values and genetic data

We used our allele frequency partitioned H-E regression framework to calculate the contribution of alleles at various frequencies to the heritability of the simulated expression levels. We corrected for population structure in both genetic and phenotypic data using the first 10 PCs from the GEUVADIS data as covariates in the analysis, as with other analyses in the paper. We subdivided the frequency space into 20 bins with approximately equal numbers of variants. For each simulated gene, we then stored the parameter values and the results of the partitioned regression in a lookup table.

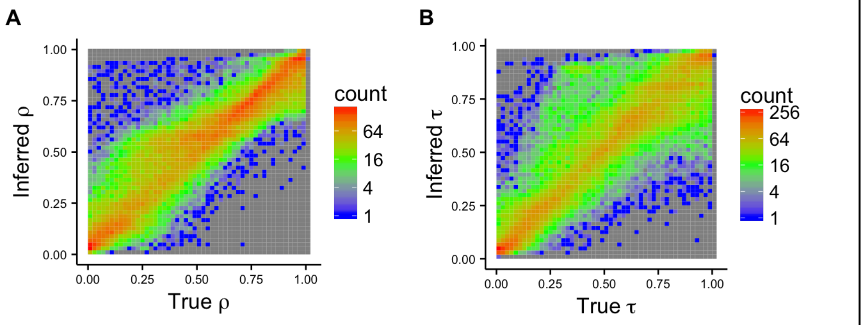


**Figure S26.** Model performance improves when parameters are large. Similar to Figure 3A-B (main text), but on the left the performance of ρ conditional on τ>0.5, and on the right the performance of τ conditional on ρ>0.5. In both cases, the estimator remains unbiased but the variance decreases.

#### 4.6 Parameter inference

To infer parameters under our evolutionary model, we used rejection sampling. We sampled each of our parameters from prior distributions (uniform from 0 to 1). We then resampled simulated genes from the lookup table described in the previous section such that mean $\rho$, $\tau$, and $\phi$ of the set match the values sampled from the prior distribution. In so doing, we assumed that there exists a beta distribution over parameter values in each gene set, and we sampled the variance over each parameter among $\rho$, $\tau$, and $\phi$ from a uniform distribution over the possible values for a beta distribution, conditional on the previously sampled mean.

As informative summary statistics for the rejection sampler, we computed the cumulative proportion of $h^2$ given by alleles in 10 frequency bins of fixed minor allele count (specifically minor allele counts of 1,2,5,10,20,60,120,180,240, and 360) for each gene and averaged the values over the whole set of 10,000 sampled genes. To determine the $h^2$ estimate for each of these frequencies, we interpolate the $h^2$ values between the 20 frequency bins from the H-E regression procedure, and use the interpolated values as summary statistics in our rejection sampling pipeline.

We repeated the above procedure 2,000,000 times, and then computed the mean squared distance between the summary statistics for each of the 2,000,000 simulations as compared to the observed data. We retained the 0.05% of the simulations, and the parameter values of mean $\tau$, mean $\rho$, and mean $\phi$, then determine the approximate posterior distribution of the parameters.

#### 4.7 Validation

To validate our parameter estimation procedure, we performed cross-validation. We masked 1,000 combinations of summary statistic and parameter combination dataset, and attempted to use the remaining 1,990,000 summary statistic and parameter combinations to infer the parameter values corresponding to the masked data.

In **Figure S25A-C**, we plot the inferred parameter value (the maximum *a posteriori* estimate) against the true parameter value for 100,000 different parameter combinations, for each of the three parameters. The tight correlation between the inferred and true parameters demonstrates that our method is accurate and relatively unbiased (note colors are plotted on a log-scale). As further validation, we computed the root mean squared deviation (RMSD) between the true and observed parameter values for both $\rho$ and $\tau$ across the space of possible $\rho$ and $\tau$ values (**Figure S25D-E**). RMSD tends to be minimized when $\tau \approx \rho$. When $\tau$ or $\rho$ is very small, our inference procedure can struggle. This is likely because when $\tau=0$, all causal alleles have the same effect size, so the parameter $\rho$ does not make sense. However, when $\rho$ and $\tau$ are both substantially greater than 0 (say >0.5), the inference is much more accurate (**Figure S26**).

As further validation of our method, in **Figure 3C** (main text), we plot out-of-sample simulations of the $h^2$ vs frequency curve, where we used each line in our inferred posterior distribution as input to a new set of resampling simulations. Since the parameter estimates are exactly those that were generated from the rejection sampling, if our procedure is robust we expect this new set of simulations to generate approximately the same $h^2$ vs. frequency curve. As expected, these out-of-sample simulations very closely track the observed data.

#### 4.8 Sufficiency of summary statistics for parameter estimation

Rejection sampling is very similar to approximate Bayesian computation (ABC)[33]. However, in ABC a linear model is imposed relating the parameter values and summary statistics, which corrects for the non-0 distance between the simulated and observed summary statistics. In rejection sampling this correction is not applied. Although the two methods are very similar, we found rejection sampling to have better empirical performance (as determined by out-of-sample simulations) for this model, and hence proceeded with this approach.

In both ABC and rejection sampling, sets of summary statistics are said to be approximately sufficient if their ability to differentiate between models is essentially equivalent to full likelihood expression. In this case, we have used the $h^2$ vs frequency curve as our set of summary statistics, but we note that in some cases (*e.g.* when $\rho$=0 or $\tau$=0) there is little this summary statistic can do to differentiate models.

We found that overall, ABC [as implemented by[34]] had very similar performance in estimating parameters to rejection sampling for this model, but also resulted in very poor fitting out-of-sample simulations when we sampled from the ABC-based posterior distribution as compared to the rejection sampling-based distribution. This difference in performance is likely due to the distinct solution spaces for $\rho$ and $\tau$ that are broadly consistent with our data. The linear model fit by ABC tends to contract all of the ABC estimates into the space in between distinct solution sets (*i.e.,* mean $\rho$ between 0.6 and 0.8, and mean $\tau$ between 0.5 and 1.0). If we were to discover and apply a fully sufficient (instead of approximately sufficient) set of summary statistics, the linear model imposed by ABC would be more appropriate because it would then not average over distinct peaks in the solution space.

#### 4.9 Predicting the population-wide frequency of causal singletons.

We find that ~25% of all $h^2$ derives from singletons in our sample of $n$=360 individuals. In **Figure 2C** (main text) we show that most of this $h^2_{\text{singletons}}$ derives from variants that are not reported in $n$>15k in gnomAD. Given our out-of-sample simulations described above (shown in **Figure 3C**, main text), we sought to predict the distribution of the true population-wide frequencies of causal variation that must contribute to $h^2_{\text{singletons}}$. We rely again on simulations. In this case, we use a feature in SFS_CODE to report the true population frequency of a segregating mutation. Using the out-of-sample draws shown in **Figure 3C** (main text), we generated the true population frequencies of the causal singletons in our $n$=360 simulations. In **Figure S27** we show that the plurality of $h^2_{\text{singletons}}$ is expected to come from variants that have true MAF <1e-5. This implies that replication will likely not be feasible for any individual variants.

7. Yang, J., Lee, S. H., Goddard, M. E. & Visscher, P. M. Genome-wide complex trait analysis (GCTA): methods, data analyses, and


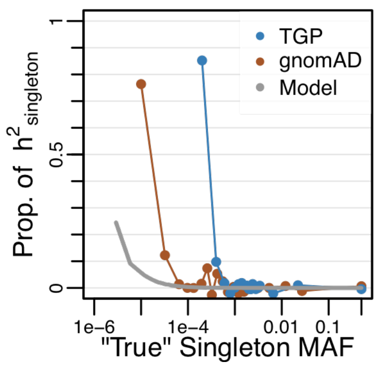


**Figure S27.** Singleton heritability is driven by extremely rare variation. gnomAD curve recreated from Figure 2B (inset, main text), and analogously partitioning variants by the MAF observed across all 2504 samples in TGP. Model shows $h^2$ as a function of "true" MAF from simulations, averaging over out-of-sample draws from the posterior distribution.

interpretations. *Methods Mol. Biol.* **1019,** 215–236 (2013).

20. Montgomery, S. B., Lappalainen, T., Gutierrez-Arcelus, M. & Dermitzakis, E. T. Rare and common regulatory variation in population-scale sequenced human genomes. *PLoS Genet.* **7,** e1002144 (2011).

27. Lohmueller, K. E. The impact of population demography and selection on the genetic architecture of complex traits. *PLoS Genet.* **10,** e1004379 (2014).

28. Simons, Y. B., Turchin, M. C., Pritchard, J. K. & Sella, G. The deleterious mutation load is insensitive to recent population history. *Nat. Genet.* **46,** 220–224 (2014).

29. Sanjak, J. S., Long, A. D. & Thornton, K. R. A Model of Compound Heterozygous, Loss-of-Function Alleles Is Broadly Consistent with Observations from Complex-Disease GWAS Datasets. *PLoS Genet.* **13,** e1006573 (2017).

# 5
